# Multi-omics analysis of virus-permissive versus hostile cellular states reveals protein networks controlling virus infection

**DOI:** 10.1101/2024.08.29.610331

**Authors:** Honglin Chen, Philip D Charles, Quan Gu, Sabrina Liberatori, David L Robertson, Massimo Palmarini, Sam J Wilson, Shabaz Mohammed, Alfredo Castello

## Abstract

The capacity of host cells to sustain or restrict virus infection is influenced by their proteome. Understanding the compendium of proteins defining cellular permissiveness is key to many questions in fundamental virology. Here, we apply a multiomic approach to determine the proteins that are associated with highly permissive, intermediate, and hostile cellular states. We observed two groups of differentially regulated genes: i) with robust changes in mRNA and protein levels, and ii) with protein/RNA discordances. Many of the latter are classified as interferon stimulated genes (ISGs) but have no reported antiviral activity. This suggests that IFN-dependent changes in mRNA levels do not imply antiviral function. Phosphoproteomics revealed an additional regulatory layer involving non-signalling proteins with altered phosphorylation. Indeed, we confirmed that several permissiveness-associated proteins with changes in abundance or phosphorylation regulate infection fitness. Altogether, our study provides a comprehensive and systematic map of the cellular alterations driving virus susceptibility.

## INTRODUCTION

The limited size of viral genomes renders viruses dependent on host cells; they often provide the molecular machineries required for virus proliferation and spread. Cellular molecules required for virus infection are globally referred to as “dependency” factors (Brass, Dykxhoorn et al. 2008, Li, Brass et al. 2009, Savidis, McDougall et al. 2016, Li, Clohisey et al. 2020). These include a wide range of proteins, including, amongst others, the receptor and co-receptors required for virus entry (Wang, Kuhn et al. 1992, Dragic, Litwin et al. 1996), the cytoskeleton to enable the transport of viral components across the cell (Arhel, Genovesio et al. 2006, Radoshitzky, Pegoraro et al. 2016), cellular co-factors that aid viral replication (Schmidt, Ganskih et al. 2023), the translation apparatus necessary for viral protein synthesis (Bushell and Sarnow 2002, Burgui, Aragon et al. 2003), the central metabolism to supply these anabolic processes with nucleotides, amino acids and energy (Ritter, Wahl et al. 2010, Cheng, Martin-Sancho et al. 2021), the glycosylation required to shield the viral glycoproteins (Pritchard, Harvey et al. 2015), and the ESCRT (endosomal sorting complexes required for transport) to release of the viral particles (Soonsawad, Xing et al. 2010, Sundquist and Krausslich 2012).

On the other hand, almost all human cell types respond to virus infection by initiating immune responses, which begin with the recognition of pathogen-associated molecular patterns (PAMPs) by cellular sensor proteins (Barbalat, Ewald et al. 2011). PAMPs include a wide range of signatures including double-stranded (ds)DNA and dsRNA, 5’-triphosphate or unmethylated caps at the end of the RNA molecules (Kato, Takeuchi et al. 2006, Sun, Wu et al. 2013). Sensing of PAMPs triggers signal transduction, which drives the expression and secretion of interferon (IFN). IFN is a family of cytokine consisting of type I, II, III that initiates the JAK-STAT signalling pathway upon binding to membrane receptors (Platanias 2005). Upon binding to its receptor, IFN triggers the expression of interferon-stimulated genes (ISGs) that facilitate virus sensing and activate mechanisms to antagonise viral replication (Sadler and Williams 2008, Schneider, Chevillotte et al. 2014). One example “effector” ISGs include the OAS-RNaseL axis: upon binding to viral RNA, the 2’-5’-oligoadenylate synthase (OAS) proteins initiate the synthesis of 2’-5’-oligoadenylate, which binds to and activates the endoribonuclease RNaseL to induce viral RNA degradation (Floyd-Smith, Slattery et al. 1981, Donovan, Dufner et al. 2013).

Permissiveness of the host cell to virus infection is collectively shaped by both host dependency and restriction factors; however, the genes and derived proteins discriminating permissive and hostile cell conditions remains under investigation. As an example, Human embryonic kidney 293 (HEK293) and its derivative HEK293T cells have distinct abilities to sustain virus infection despite their close lineage. This is evidenced by studies showing that HEK293T cells can produce higher titre of murine stem cell virus and human immunodeficiency virus type 1 (HIV-1) (Li, Kao et al. 2012), and HIV-1-based lentivirus particles (Merten, Charrier et al. 2011, Ferreira, Sumner et al. 2020). HEK293T also produce a higher level of infective adenovirus particles (Park, Lim et al. 2006, Bae, Marino et al. 2020, De, Cram et al. 2023), Epstein-Barr virus (Chen, Sathiyamoorthy et al. 2018), and Influenza A virus under co-culture conditions (Milian, Julien et al. 2017), and a higher capacity to sustain SARS-CoV-2 replication (Harcourt, Tamin et al. 2020, Modrof, Kerschbaum et al. 2020). HEK293T originates from stable transfection of HEK293 with Simian virus 40 large tumour antigen (SV40-LT) (Rio, Clark et al. 1985); however, its different level of permissiveness to viruses cannot be solely attributed to expression of SV40-LT, because its depletion has a limited effect on virus yield (Bae, Marino et al. 2020, Ferreira, Sumner et al. 2020), and SV40-LT has multifaceted impacts on innate immunity (Swaminathan, Rajan et al. 1996, Forero, Giacobbi et al. 2014, Lau, Gray et al. 2015, Reus, Trivino-Soto et al. 2020). SLF11 is more abundant in HEK293 than in HEK293T cells, and it was proposed to inhibit HIV-1 by controlling the host aminoacyl-tRNA pool (Li, Kao et al. 2012). However, following experiments showed that SLF11 cannot fully explain the differential ability of HEK293 and HEK293T to sustain HIV-1 infection, and this host factor does not regulate other viruses (Li, Kao et al. 2012). The genetic variations revealed by a large-scale genome sequencing study comparing HEK cell lines also fails to explain the distinct permissiveness of HEK293 and HEK293T (Lin, Boone et al. 2014). The determinants of virus permissiveness between these closely related cell lines thus remains largely unknown.

HEK293 and HEK293T cell lines provide a model system to study determinants of virus permissiveness due to their differential capacity to sustain infection despite having closely related genomes (Boone, Lin et al. 2014). In this study, we employed an integrated omics analysis to uncover the scope of cellular proteins that define virus permissiveness across multiple cellular states, including the HEK293T cells (highly permissive), steady-state HEK293 cells (intermediate), and HEK293 upon stimulation of IFN-α (hostile). Our results revealed a subset of antiviral factors that are globally depleted in HEK293T cells while anabolic pathways are upregulated, creating an ideal environment for virus replication. In addition, our transcript- and proteomics data depicted a temporal gene expression regulation in the IFN-α response. This analysis pinpointed a group of antiviral factors that is robustly and reproducibly upregulated in response to IFN-α. Moreover, we observe another group of genes that despite being upregulated at the RNA level, do not show changes at the protein level, which suggests IFN-α-induced transcriptional noise with limited effects at regulating infection. Moreover, our phosphoproteomic analysis uncovered extensive modulation of phosphorylation among non-signalling proteins in response to IFN-α. Through integrated analysis of omics data, we identified novel regulators of virus infection, and provided experimental validations on their regulatory roles against Sindbis virus (SINV) and HIV-1.

## RESULTS

### Widespread proteome differences between HEK293 and HEK293T

Previous work reported that infection of multiple viruses, including HIV-1, proceeds more efficiently in HEK293T cells than in HEK293 cells. To further investigate these results, we infected HEK293 and HEK293T with an HIV-1_Gag-mCherry_ replicon (suppl.f1a) pseudotyped with the glycoprotein of vesicular stomatitis virus (VSV-G), and viral gene expression was measured using mCherry as a proxy as in (Garcia-Moreno, Noerenberg et al. 2019). We observed a substantially higher HIV-1_Gag-mCherry_ derived fluorescent signal in HEK293T than in HEK293 cells throughout the course of infection (fig. 1a, top). Consistently, Gag and p24 abundance was higher in HEK293T than in HEK293 cells for both HIV-1_Gag-mCherry_ or HIV-1_Nef-mCherry_ (fig. 1a, bottom; suppl.f1b, left) (Garcia-Moreno, Truman et al. 2023). To generalise these results, we infected HEK293 and HEK293T with SINV, a vector-borne RNA virus within the *Togaviridae* family (SINV_mCherry_; suppl.f1a) (Garcia-Moreno, Noerenberg et al. 2019). Analysis of mCherry and capsid abundance confirmed that SINV infects more efficiently HEK293T than HEK293 (fig. 1a; suppl.f1b). Altogether, these results and those reported previously (Park, Lim et al. 2006, Merten, Charrier et al. 2011, Li, Kao et al. 2012, Milian, Julien et al. 2017, Chen, Sathiyamoorthy et al. 2018, Bae, Marino et al. 2020, Ferreira, Sumner et al. 2020, Modrof, Kerschbaum et al. 2020, De, Cram et al. 2023) support that HEK293T are more permissive to virus infection than HEK293.

**Figure 1.**
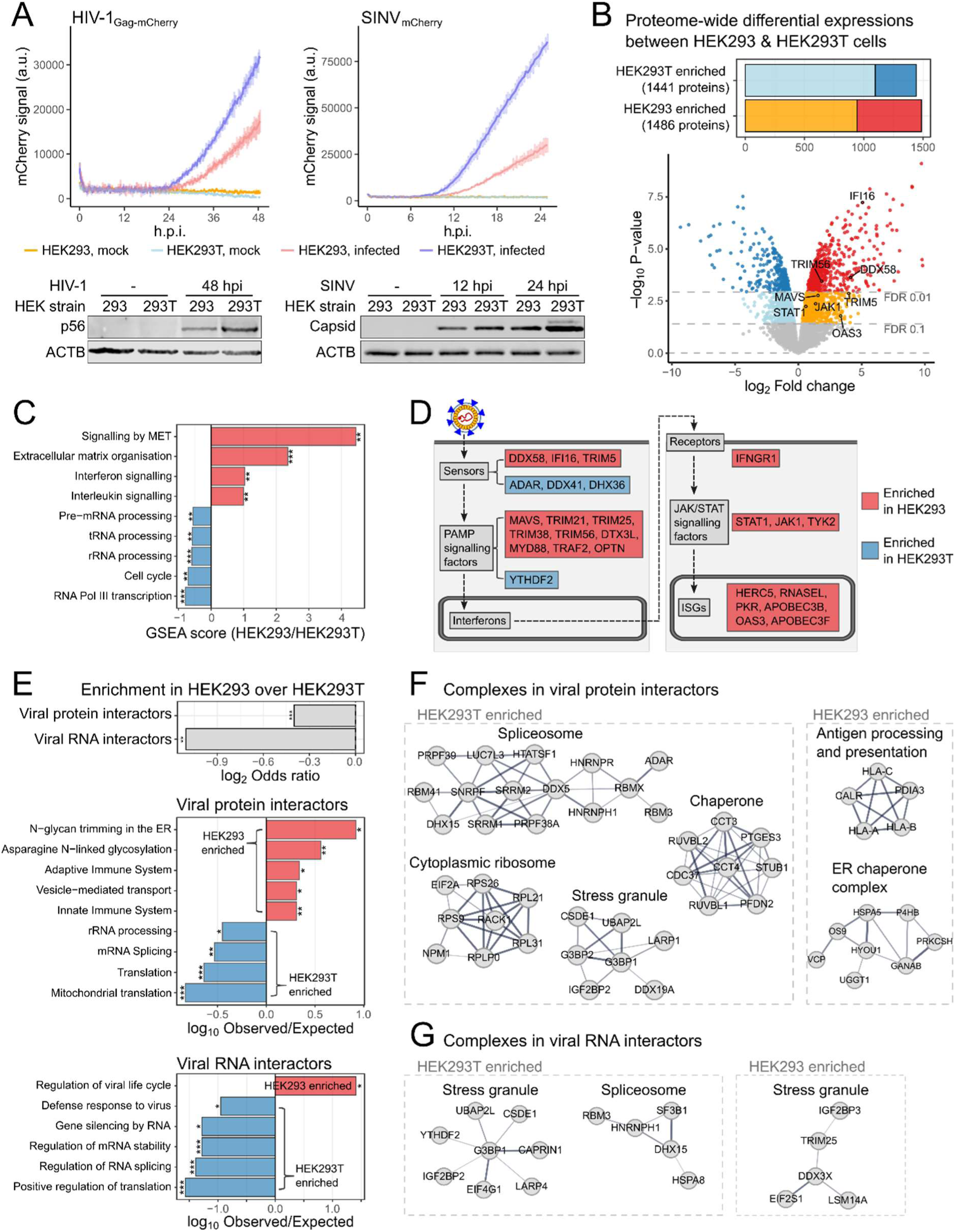
Host factors and processes underlying different permissiveness between HEK293 and HEK293T cells. A) Infection fitness of HIV-1_Gag-mCherry_ (left) and SINV_mCherry_ (right) in HEK293 and HEK293T cells. Fluorescence signals were measured every 15 min in culture condition using a plate reader (top, n=3). Western blottings were performed using HIV-1_Nef-mCherry_ (left) and SINV_mCherry_ (right) with antibodies of viral capsid proteins (bottom, multiplicity of infection (MOI) =1). B) Result of quantitative proteomic comparison between HEK293 and HEK293T. Bar plot summarised numbers of proteins with FDR<0.1 in each cell line (top). Volcano plot showed log2 fold change and significance (p-value) of each protein between HEK293 and HEK293T cells (bottom, suppl.tab1). C) Differentially enriched *Reactome* pathways in Gene set enrichment analysis (GSEA). P-values of enrichments were * labelled (**: p<0.01, ***: p<0.001). D) Schematic of innate immune response against virus with key processes labelled in grey boxes. Differentially enriched genes in each process were labelled red (HEK293-enriched) or blue (HEK293T-enriched). E) Enrichments of viral protein interactors and viral RNA interactors between HEK293 and HEK293T cells (top), and Gene ontology – Biological processes (GO-BP) terms enrichments of viral protein interactors (mid) viral RNA interactors (bottom) in HEK293 (red) and HEK293T (blue) cells. P-value represented by * label (*: p<0.05, **: p<0.01, ***: p<0.001) F) Protein-protein interaction complexes among viral protein interactors differentially enriched between HEK293 and HEK293T cells. G) As in (F) but for viral RNA interactors.

To explore factors for the differential ability of HEK293 and HEK293T cells to sustain infection, we conducted whole-cell proteome profiling with stable isotope labelling with amino acid in culture (SILAC) and offline high-pH RP fractionation, achieving over 99% isotope incorporation (suppl.f1c)(Ong, Blagoev et al. 2002, Di Palma, Hennrich et al. 2012). We identified and quantified 8553 and 7558 proteins, respectively (suppl.1d,e). Despite the shared genetic background of the two lines, ∼3k proteins exhibited differential abundance at 10% false discovery rate (FDR), with 1,486 and 1,441 proteins being enriched in HEK293 and HEK293T cells, respectively (fig. 1b). Gene set enrichment analysis revealed a prevalence of interferon signalling pathway in HEK293 cells over HEK293T cells (fig. 1c). Further investigation identified proteins involved in virtually all processes of the antiviral response that are more abundant in HEK293 cells than in HEK239T (fig. 1d). These included: i) intracellular sensors such as RIG-I (DDX58) and IFI16 (Pichlmair, Schulz et al. 2006, Garcia-Moreno, Noerenberg et al. 2019, Kim, Arcos et al. 2020), ii) effectors such as TRIM5, PKR, OAS3, and RNASEL (Stremlau, Owens et al. 2004, Garcia, Meurs et al. 2007, Silverman 2007), iii) signal transducers such as MAVS and TRAF2 (Hou, Sun et al. 2011), iii) receptor signalling factors such as MYD88 and the tyrosine kinases JAK1 and TYK2 (Silvennoinen, Ihle et al. 1993, Dunne, Ejdeback et al. 2003), and iv) the transcription factor STAT1 (Darnell 1997).

The widespread depletion of innate immunity proteins in HEK293T cells implies a globally dysfunctional antiviral response that cannot be ascribed to individual proteins (Li, Kao et al. 2012). To verify this, we treated both cell lines with IFN-α and measured the level of STAT1 Y701 phosphorylation (suppl.f1f). Interestingly, blotting results showed that both cell lines respond to IFN-α stimulation, but with a different strength (suppl.f1g), and saturation dose (suppl.f1h). Differences in strength and saturation point of the IFN-α stimulation are presumably derived from the distinct stoichiometry of the IFN receptors and signalling proteins in the two lines.

Beyond innate immunity, hundreds of proteins involved in other cellular processes were also differentially expressed in HEK293 and HEK293T cells. To identify regulators of infection that are outside the scope of well-characterised ISGs, we cross-referenced our dataset with a manually compiled and curated list of viral protein and RNA interactors from previous publications (suppl.f1I) (Jager, Cimermancic et al. 2012, Pichlmair, Kandasamy et al. 2012, Shah, Link et al. 2018, Gordon, Jang et al. 2020, Iselin, Palmalux et al. 2022). Viral protein and RNA interactors in HEK293T are enriched in gene ontology (GO) terms related to RNA metabolic processes that are typically classified as dependency factors, including the translation and splicing apparatus and RNA stability factors (fig. 1e-f). Conversely, the viral protein interactors in HEK293 cells are involved in immune response and protein N-glycosylation (fig. 1e, mid), including MHC-I and MHC-I peptide loading complex (fig. 1f, right). Notably, non-core stress granule components showed different abundance in HEK293 and HEK293T (fig. 1g). YTHDF2 is enriched in HEK293T and was reported to inhibit innate immune response through RNA methylation-dependent degradation (Winkler, Gillis et al. 2019). By contrast TRIM25 and LSM14A are associated with antiviral functions and are enriched in HEK293 (Gack, Shin et al. 2007, Li, Chen et al. 2012). Altogether, our data support a model in which the distinct permissiveness of HEK293 and HEK293T is derived from complex proteome differences involving a wide range of antiviral and dependency factors.

### The proteome landscape of HEK293 cells after IFN-α stimulation

We next extended the comparative proteomics analysis to include a virus-hostile cellular state. We focused on IFN-α treated HEK293 given that HEK293T cells possess a compromised antiviral activity even in presence of IFN-α (fig. 2f; suppl.f2a). We observed a mild but significant inhibition of SINV gene expression just with 10 min of IFN-α treatment prior to infection, which increased in magnitude when the pre-treatment was extended to 4 and 20 h (suppl.f2b,c). To profile the proteome responses, SILAC-labelled HEK293 cells were treated with mock or 200 U/ml IFN-α for 10 min, 4 h, or 20 h (fig. 2a). Proteome changes are expected to require hours, while posttranslational modifications (PTMs) can occur within minutes. Therefore, we used the 10 min and 4h samples for phosphoproteomics and the 4 and 20h samples for deep proteome analyses (fig. 2a).

**Figure 2.**
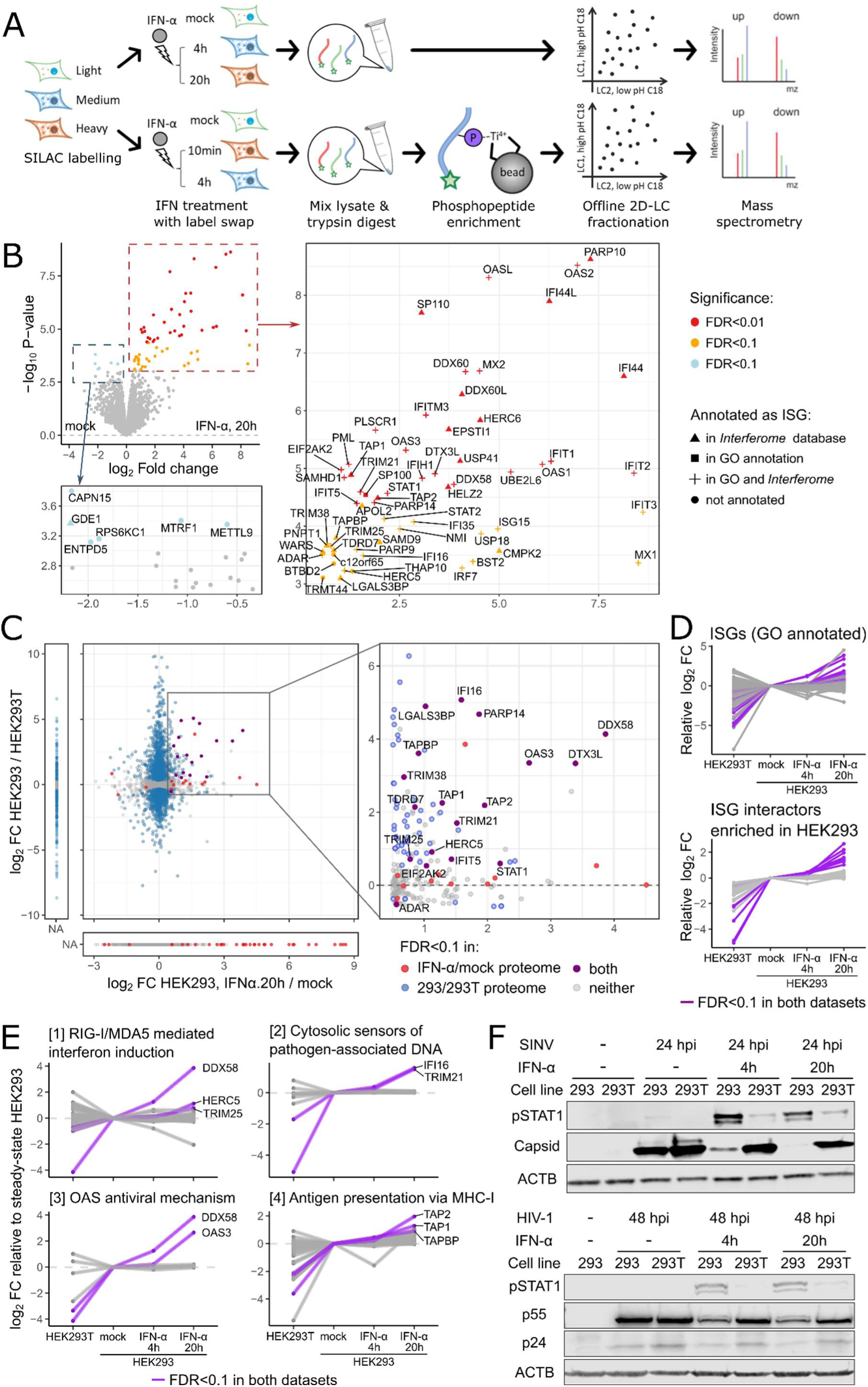
Proteome landscape of HEK293 cells after IFN-α stimulation. A) Schematic workflow of proteome and phosphoproteome profiling of IFN-α stimulated HEK293 cells. SILAC samples were mixed with label swap between replicates. B) Volcano plot with text label showing proteins with FDR<0.1 after 20h IFN-α treatment (suppl.tab2). Proteins related to IFN-α were drawn in different shapes depending on source of annotations – triangle (Δ): annotated in microarray (MA) studies in database *Interferome*; square (□): annotated in GO-BP terms; plus (+): annotated in both MA studies and GO-BP terms; circle: no IFN-α related annotation in either source. C) Scatter plot comparing log2 fold changes quantified in IFN-α/mock proteome and HEK293/HEK293T proteome for each protein. D) Parallel coordinate plots showing protein expression patterns across cellular states. Proteins with FDR<0.1 in both IFN-α/mock and HEK293/HEK293T proteomes were colour labelled. E) As in (D) but showing proteins in pathways of interests. F) Western blotting with antibodies of STAT1-pTyr701 (pSTAT1) and viral capsid proteins. Cells were infected at a MOI=1 with SINV_mCherry_ (top) or HIV-1_Nef-mCherry_ (bottom).

Deep proteome profiling identified and quantified 10,154 and 9,131 proteins (suppl.f2d-f). To our surprise, only 66 proteins showed significant changes at 20 hpt under a cut-off of FDR<0.1 (fig. 2b). This is a small number given that 7,350 human genes are reported to respond to type I IFN in micro array (MA)-based studies in the *Interferome* database (Rusinova, Forster et al. 2013). Among the proteins with significant upregulation upon IFN-α treatment, 37 were classified as ISGs by GO annotation, 55 by MA, and 36 by both (fig. 2b). Four proteins were not linked to IFN-α response according to the referenced sources. By contrast, none of the IFN-α downregulated proteins were related to innate immunity except for GDE1 that was detected in one MA study (Henig, Avidan et al. 2013). The depth and precision of this proteome dataset also allowed depiction of temporal changes across early (4 hours post treatment, hpt), and late (20 hpt) timepoints. Quantified proteins were divided into 8 groups based on their temporal profile (suppl.2g). Groups showing quick and continuous or late induction upon IFN-α stimulation were enriched in antiviral response pathways, including RIG-I/MDA signalling, IFN-α signalling, and ISG15 mediated antiviral mechanisms (suppl.f2h). Several known antiviral proteins were present in a third group showing quick induction followed by plateau, including the transcription factor IRF9 and RIG-I (DHX58) (suppl.f2g, middle-right). Most proteins in the other groups have no known role in innate immunity.

To search for factors that modulate virus permissiveness across different cellular states, we compared proteomic data from permissive HEK293T cells, steady-state HEK293 cells, and hostile IFN-α stimulated HEK293 cells (fig. 2c). We found 17 proteins enriched in virus-restrictive IFN-α stimulated HEK293 cells and depleted in virus-permissive HEK293T (fig. 2c). Thirteen of these are functionally well annotated ISGs, suggesting that these may represent a pivotal antiviral network defining cell susceptibility to virus infection. These ISGs displayed a gradual increase in protein abundance as the cellular state becomes more virus-restrictive (fig. 2d, top). Pathway analysis determined that they are involved in viral RNA and DNA sensing, and 2’-5’-oligoadenylate synthesis (fig. 2e). Moreover, we observed several proteins described as interactors of ISGs that also increased in abundance as cells become more restrictive (fig. 2d, bottom) (Hubel, Urban et al. 2019). These ISG interactors include LGALS3BP that is as a scaffold protein for PAMP signalling with broad spectrum antiviral roles (Xu, Xia et al. 2019), and TAP1/2 are antigen presentation factors that were linked to innate immunity in functional studies (Tanaka, Hosokawa et al. 2011, Blees, Januliene et al. 2017); however, their roles in IFN response are not well understood.

### Transcriptome and proteome profiling reveals discordances between RNA and protein levels during IFN-α response

To investigate the relationship between transcript and protein presence, we performed RNA-seq with IFN-α stimulated and untreated HEK293 cells. Our results revealed that 126 and 419 genes exhibited significant differential expression at 4 and 20 hpt, respectively (fig. 3a, suppl.f3a). Comparing this RNA-seq result with similar studies in different human cell lines, we found that numbers of significantly up- and downregulated genes vary substantially across datasets (suppl.f3b), which can reflect differences in cell types and experimental conditions (Shaw, Hughes et al. 2017, Colli, Ramos-Rodriguez et al. 2020). Despite these quantitative differences, we observed high correlations among genes that were commonly upregulated across cell types (R = 0.63 and 0.73, suppl.f3c). In addition, a large proportion of commonly upregulated genes are GO-annotated ISGs (41%), while less than 4% of genes that exhibited cell type-specific regulations are linked to IFN response (suppl.f3d). These observations support that IFN-α responses in HEK293 cells are representative of ISGs with robust upregulation across cell types.

**Figure 3.**
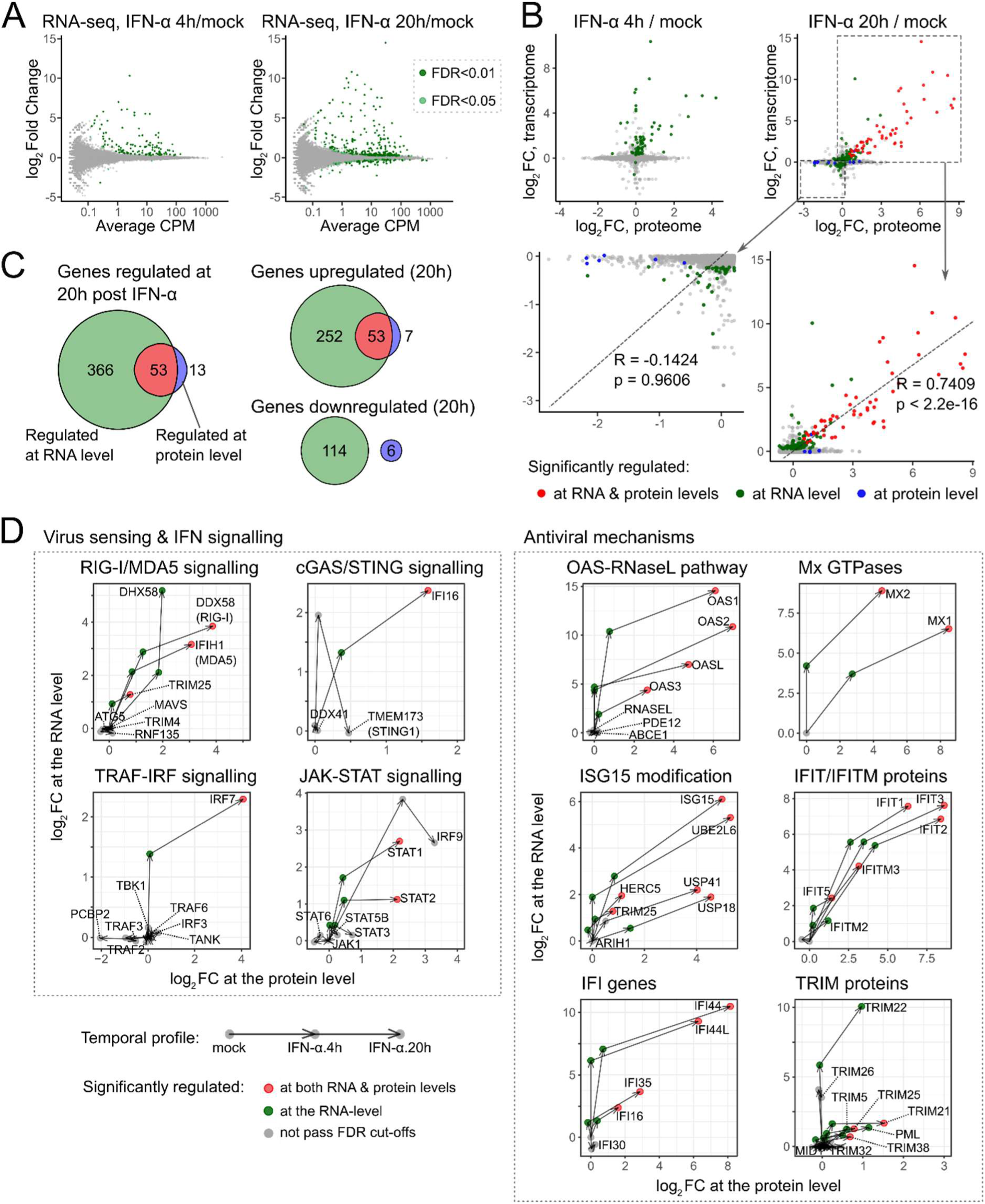
Transcriptome analysis of HEK293 cells after IFN-α stimulation. A) MA plots of IFN-α induced transcriptome changes at 4 hpt (left) and 20 hpt (right) in HEK293 cells. Gene expression was measured by CPM (counts per million reads), referring to the number of reads mapped to transcript scaled by the number of sequencing reads (suppl.tab3). B) Scatter plots comparing log2 fold changes in transcriptome and proteome after IFN-α stimulation. Only genes quantified in both datasets were plotted. C) Venn diagrams comparing numbers of IFN-α induced significant changes at 20 hpt at RNA and protein levels. D) Vector plots visualising temporal expression profiles after IFN-α stimulation at RNA and protein levels.

We next compared our transcriptome and proteome datasets to assess correlations between RNA and protein levels at different timepoints. When focusing exclusively on genes upregulated at the RNA and protein level, we observed a positive trend at both 4 and 20 hpt (R = 0.74, fig. 3b bottom-right). When equal weighting was applied to all genes and also considered genes only quantified by one method, we found that 366 out of the 419 genes regulated at the RNA level had no corresponding changes at protein level (fig. 3c, left). We observed good correlation for between RNA and protein for 53 upregulated genes, mostly ISGs. Surprisingly, we noticed a striking lack correlation between RNA and protein for downregulated genes (fig. 3b-c). This discordance was also observed when using other published RNA-seq experiments (suppl.f3b) (Shaw, Hughes et al. 2017, Colli, Ramos-Rodriguez et al. 2020). We next assessed if the observed RNA-protein discordance was caused by different detection limits between RNA-seq and proteomics. Interestingly, ∼40% of the genes with changes only at the RNA level were also quantified by our proteomic analysis (suppl.f3e). This indicates that a substantial proportion of RNA-level changes are not reflected at the protein level, among which we observed a series of well-studied ISGs such as IRF1/2, IFI6/27 (suppl.f3f) (Harada, Fujita et al. 1989, Meyer, Kwon et al. 2015, Xue, Yang et al. 2016).

To better understand the gene expression kinetics during IFN-α response, we visualised temporal kinetics profiles with vector plots connecting RNA- and protein-level changes at different timepoints on the same coordinate map (fig. 3d). Most genes within the strongest upregulated pathways displayed similar temporal kinetics. These genes first increase at the RNA level, followed by an upregulation of both RNA and protein at 20 hpt (fig. 3d). These profiles agree with a transcription/translation driven gene expression model, that requires time to transform the transcriptional response into protein (Schwanhausser, Busse et al. 2011). This temporal gap should be considered when analysing IFN responses with RNA-centric methods.

### Protein-level IFN-α response has higher correlation with characterised antiviral functions

We then assessed functional annotations for genes that display discordance between RNA and protein levels. Indeed, we noticed that genes regulated at both RNA and protein level were heavily annotated with IFN-related GO terms, while the opposite was observed with genes exhibiting changes only at RNA or protein level (fig. 4a). We next inspected a series of large-scale functional screenings that assessed antiviral activities of close to 400 ISGs against 17 RNA viruses (fig. 4b) (Schoggins, Wilson et al. 2011, Schoggins, MacDuff et al. 2014). We found a prevalence of ISG with experimentally determined antiviral activity within the group of proteins upregulated at both RNA and protein levels (fig. 4c). These ISG screenings also highlighted a group of broad-acting ISGs with inhibitory effects for multiple viruses spanning several families (Schoggins, Wilson et al. 2011). These ISGs are pivotal in the antiviral response, and include well-characterised sensors (RIG-I/DDX58, MDA5/IFIH1), transcription factors (IRF and STAT families), and effectors (OAS, MX, and IFI families). Strikingly, these broad-spectrum ISGs are more prevalent within the genes with correlation at the protein and RNA levels than in the discordant genes (fig. 4d,e; suppl.f4b). Another large-scale study defined the “core ISG set”, which included genes with robust stimulation in response to type-I IFN across mammals (Shaw, Hughes et al. 2017). Again, the genes with robust upregulation at the RNA and protein levels are enriched in “core-ISGs” when compared with the discordant genes (suppl.f4a). Altogether, our results pinpoint a strong correlation between pivotal antiviral factors and the robust response to IFN stimulation involving both RNA and protein.

**Figure 4.**
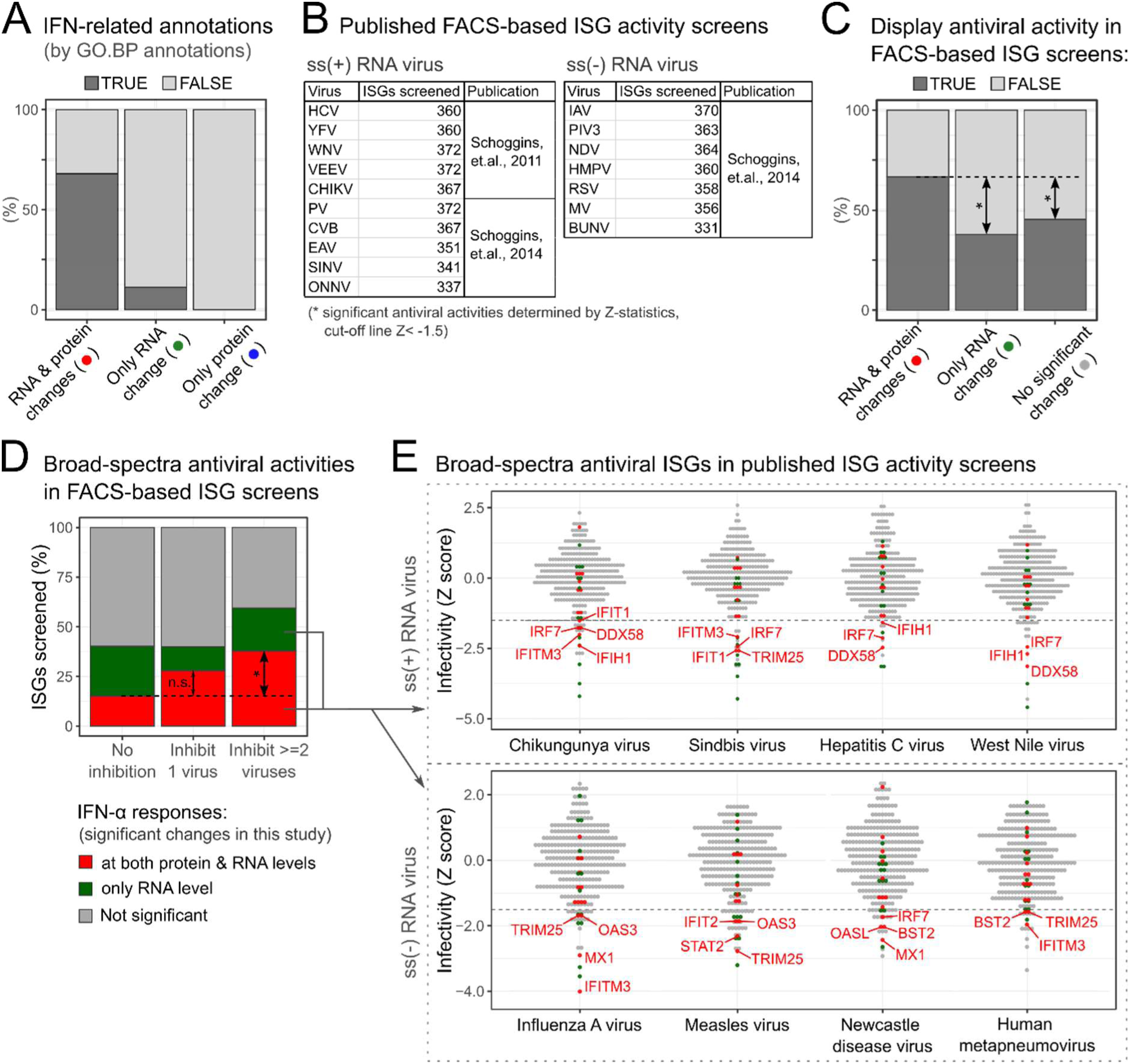
Correlation analysis with characterised antiviral functions. A) Proportions of GO-BP annotated ISGs among genes regulated by IFN-α in HEK293 cells at RNA/protein levels. B) Tables summarising publications, virus species, and numbers of ISGs assessed in fluorescence-activated cell sorting (FACS) based antiviral activity screenings. C) Proportions of FACS screening-verified antiviral ISGs among genes regulated by IFN-α in HEK293 cells at RNA/protein levels. D) Proportions of broad-spectrum antiviral ISGs among genes regulated by IFN-α in HEK293 cells. ISG activities in FACS screening results were grouped by the number of viruses they can inhibit: no inhibition, inhibit 1 virus (virus-specific), and inhibit ≥ 2 viruses (broad-spectra). E) Dot plots of FACS screening results listed in (B). Genes with broad-spectrum antiviral activities were colour labelled based on IFN-α response at RNA/protein levels in HEK293 cells. ISGs with Z < −1.5 for each virus were text labelled.

We next focused on the differentially regulated genes with no IFN-related GO annotation. We noticed that many of these poorly understood genes clustered together in a protein-protein interaction network, suggesting functional interconnections (suppl.f4c). Several highly connected hubs (nodes with ≥ 5 connections) in this network included CMPK2 and IFI44, which have recently been reported as antiviral factors (Busse, Habgood-Coote et al. 2020, Lai, Wu et al. 2021). HERC6 is another network hub, which is a catalytically inactive homolog of the known antiviral factor HERC5 (Wong, Pung et al. 2006) with no yet described function in immunity. Interestingly, genes regulated only at the RNA level were enriched in apoptosis factors (suppl.f4d). Proteome data confirmed that many key regulators of apoptosis were quantified but had no significant changes at the protein level upon IFN-α stimulation, as it is the case of the pivotal apoptotic proteases CASP4, 7 and 8 (Fan, Han et al. 2005) (suppl.f4e). These genes showed a distinct induction profile when compared with ISGs, with low to no increase at 4hpt and a strong upregulation at 20hpt (suppl.f4f). A plausible explanation for these results is that ISGs represent a first transcriptional wave, followed a second wave including the apoptotic programme if the IFN-α stimulation is prologued in time. The delayed production of mRNAs encoding pro-apoptotic factors would inevitably result in the accumulation of these proteins at later times post exposure, which were not covered in our proteomic analysis.

### Phosphoproteomics links IFN-α response to RNA-binding proteins

To assess if post-translational modulations contribute to define cell permissiveness, a deep phosphoproteome analysis was conducted in cells treated with IFN-α for 10 min or 4h (fig. 2a). We identified 23,377 phosphosites in 3,942 proteins (suppl.tab3 and suppl.f5a-c). Our analysis identified 145 phosphosites with altered abundance, and an additional 285 sites with “ON-OFF” changes (fig. 5a, suppl.f5d). When mapped to functional annotation databases (Hornbeck, Kornhauser et al. 2012), we found that only 10.3% of IFN-α regulated sites have supporting regulatory functions, 1.5% have known roles in signalling pathways, and 3.6% map to known ISGs (fig. 5b). A similar proportion of functional annotation was also observed for the total set of identified phosphopeptides, irrespective of IFN-α regulation (suppl.f5e). These results are consistent with a published meta-analysis suggesting that the vast majority of recorded phosphosites have no reported function (Needham, Parker et al. 2019). Strikingly, we detected changes in key nodes of the innate immune signalling, including the nuclear body associated protein PML (El Bougrini, Dianoux et al. 2011) and ANKRD17 (Wang, Tong et al. 2012) (suppl.f5g). We further investigated processes underlying the observed phosphosite regulations using kinase-substrate enrichment analysis (KSEA) (Lachmann and Ma’ayan 2009). We mapped 11,305 kinase-substrate connections across 390 kinases and 4,698 sites, uncovering 18 kinases with significantly altered activities (suppl.f5h, suppl.tab4). A few of these kinases are linked to immune response, including IKBKE that activates IRF3 and STAT proteins (Gu, Fullam et al. 2013), and WNK kinases (Pichlmair, Kandasamy et al. 2012).

**Figure 5.**
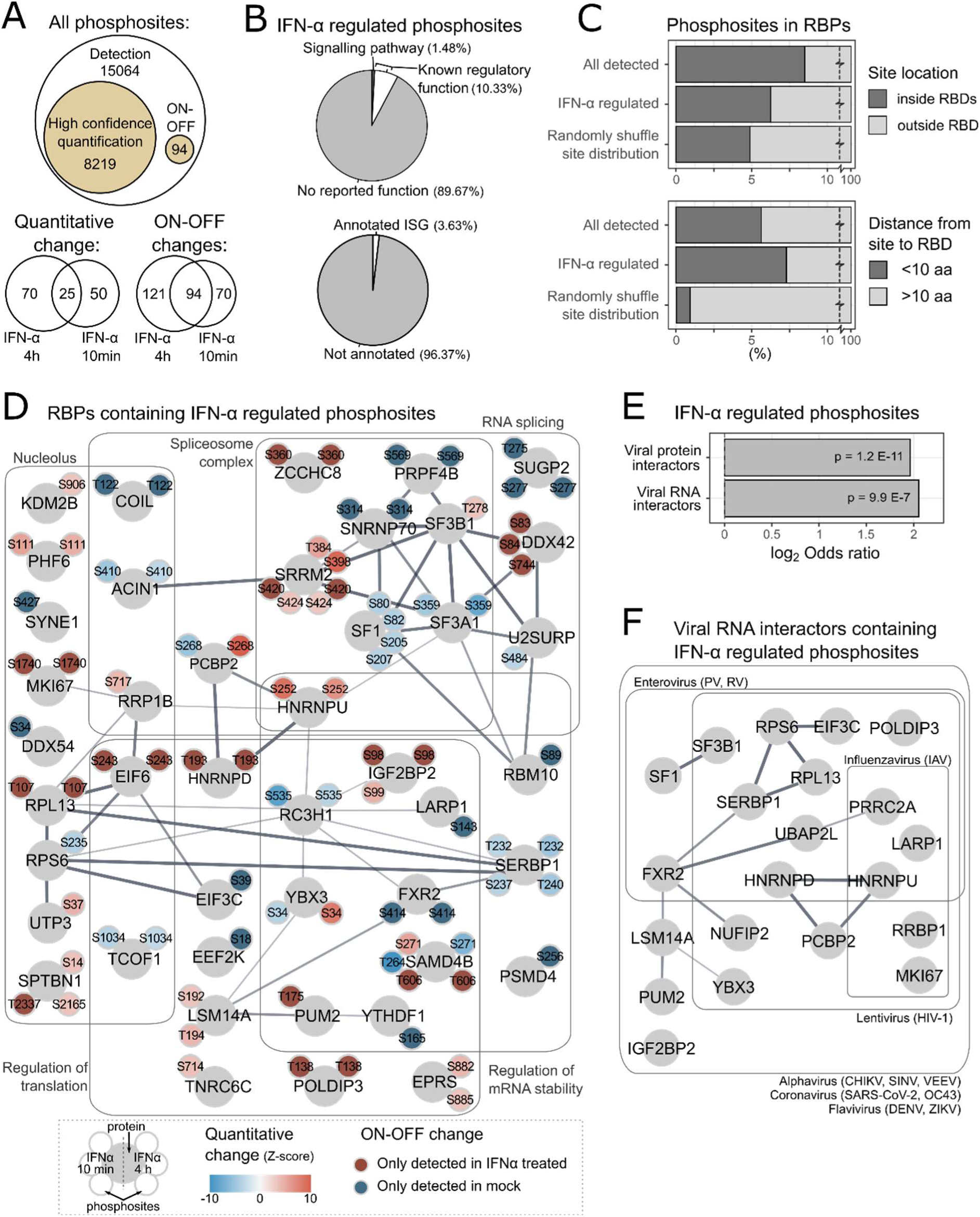
Phosphoproteome analysis of HEK293 cells after IFN-α stimulation. A) Venn diagrams summarising numbers of phosphosites identified, quantified, and IFN-α regulated in phosphoproteomic result that combined data from two instrument configurations (suppl.tab4) B) Proportions of functional annotations among IFN-α regulated sites (top) and IFN-related GO-BP annotations among phosphoproteins that contain IFN-α regulated phosphosites (bottom). C) Proportions of phosphosites that locate inside annotated RNA binding domains (RBDs; top), or locate within 10 amino acids before or after RBDs (bottom). D) Protein-protein interaction network of RNA binding proteins (RBPs) that contain IFN-α regulated phosphosites. Positions of phosphosites were labelled with small circles adjacent to each RBP. E) Enrichments of viral protein interactors and viral RNA interactors among proteins that have IFN-α regulated phosphosites. F) Protein-protein interaction network of viral RNA interactors that have IFN-α regulated phosphosites. Virus family and species that each protein interacts with were text labelled.

RNA-binding proteins (RBPs) are central to virus infection acting as dependency and antiviral factors (Castello, Álvarez et al. 2024). Critically, their RNA-binding domains (RBDs) are enriched in PTMs, which can modulate the interaction with RNA (Castello, Fischer et al. 2016). Here we assessed if phosphorylation could regulate cellular RBPs during IFN-α response. Phosphosites detected in our dataset were indeed prevalent inside known RBDs, although this enrichment was more moderate when considering only the IFN-α regulated phosphosites (fig. 5c top). Strikingly, IFN-α modulated phosphosites became strongly enriched when RBDs were considering the 10 amino acids before and after the RBD (fig. 5c bottom, suppl.f5i). The proximity between IFN-α regulated sites and RBDs indicates that phosphorylation-dependent regulation of RBP could play a role in IFN-α response. Protein-protein interaction network revealed that RBPs containing IFN-α regulated sites were widely involved in gene expression control, including regulation of splicing, translation, and RNA degradation (fig. 5d). Cross-referencing analysis confirmed significant enrichments of IFN-α regulated phosphoproteins among known interactors of viral RNA and proteins, with several of them being capable to bind the RNA of viruses from different species and families (fig. 5e-f). One of these broad-spectrum interactors, HNRNPD, was phosphorylated after IFN-α treatment at T193, which localises to its RBD (suppl.f5j); while POLDIP3 was phosphorylated at T138 close to its non-canonical RBD (suppl.f5j). These observations implied that interactions with viral RNA could be subject to phosphorylation-dependent regulation during IFN-α response.

### Uncovering new regulators of virus infection

To test if additional antiviral proteins exist beyond known ISGs, we selected 15 candidates with differential abundance or phosphorylation status in IFN-α, prioritizing proteins with little or no previous implications in immunity (suppl.tab5). These candidates were expressed in a doxycycline-dependent manner fused to eGFP and used to challenge HIV-1_Gag-mCherry_ and SINV_mcherry_ infection (suppl.f6a). We used TRIM25-eGFP and eGFP as positive and negative controls, respectively (Garcia-Moreno, Noerenberg et al. 2019). A significant decrease of red fluorescence signal derived from HIV-1_Gag-mCherry_ was observed for 6 out of 9 tested genes (fig. 6a; suppl.f6b,c). Five of these phenotypes were confirmed by analysis of Gag/p24 expression by Western blotting (fig. 6a; suppl.f6d). Interestingly, both E2 ubiquitin ligases UBE2L6 and UBE2J2 showed inhibitory effect against HIV-1 (fig. 6a, suppl.f6b). While UBE2L6 is a known ISG15 ligase in innate immune response (Zhao, Beaudenon et al. 2004), UBE2J2 has no documented role in HIV-1 infection. However, UBE2J2 reduced to nearly half the expression of p24 (fig. 6a, suppl.f6d). Nuclear-localised RBP HNRNPD harbour an IFN-α induced phosphosite and, interestingly, our data showed that its overexpression can suppress HIV-1 replication (fig. 6a).

**Figure 6.**
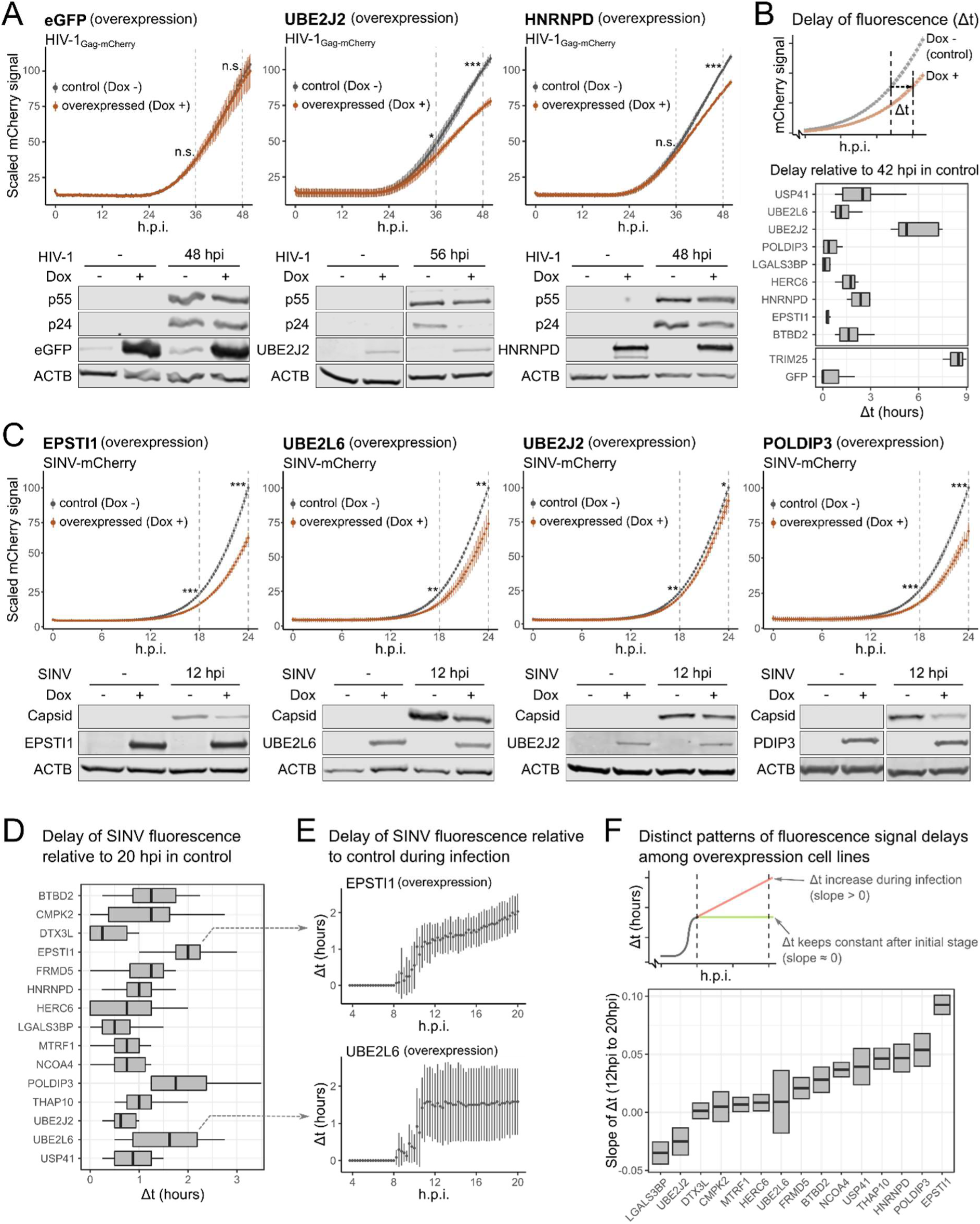
Virus-inhibitory effects against SINV and HIV-1 in stable cell lines expressing candidate genes. A) Infection fitness of HIV-1_Gag-mCherry_ in HEK293 inducible stable lines expressing candidate genes with eGFP fusion (top, MOI = 1, n = 3). Fluorescence signals were measured as in (fig. 1A). Western blotting analysis were performed with anti-eGFP and anti-p24 (bottom, MOI = 1). B) Schematic of fluorescence signal delay analysis that assess delays in virus gene expression caused by expression of candidate gene (top), and delay analysis results at 42 hpi for all candidates assessed with HIV-1_Gag-mCherry_ (bottom). C) As in (A) but with SINV-mCherry for fluorescence assay and Western blots. D) As in (B) but with SINV-mCherry at 20 hpi. E) Fluorescence signal delay analysis from 4 to 20 hpi for EPSTI1 and UBE2L6 overexpression cells. F) Schematic of delay pattern analysis that assess slope of delays (top), and comparison of delay patterns between 8 and 20 hpi for all candidates assessed with SINV-mCherry (bottom).

To quantitatively evaluate HIV-1 inhibitory effects, we assessed the delay in viral gene expression caused by the overexpression of the candidates in our near real-time plate reader assay (fig. f6b, top). We first determined the mCherry fluorescence signal at 42hpi in the control cells, and then assessed the time such signal levels were achieved in the different overexpression lines, which we refer to as “delay” or Δt. Untagged eGFP expression caused a delay relative to the uninduced line that was negligible, similarly to EPSTI1, LGALS3BP and POLDIP3 expression. However, expression of several eGFP-fused candidates led to delays ranging from 2 to 5h, with UBE2J2 causing a delay comparable to TRIM25 (fig. 6b, bottom). We then extended the delay analysis to the 12∼42 hpi window. GFP control and non-inhibitory candidates showed near a 0h delay throughout this time window, while fusion proteins with anti-HIV-1 effects had continuous increases in Δt after 24 hpi (suppl.f6e).

We next extended the study to a virus from a different family, SINV_mCherry_. Interestingly, 13 out of 15 tested fusion proteins caused a statistically significant decrease in mCherry fluorescent signal compared to the uninduced cells (fig. 6c, suppl.f6f,g). The inhibitory phenotypes of 5 fusion proteins were confirmed by Western blots (fig. 6c, suppl.f6h). EPSTI1, which showed no effect on HIV-1, caused the strongest anti-SINV phenotype (fig. 6c). Interestingly, the antiviral activity of EPSTI1 was reported for other positive sense, single stranded RNA virus, hepatitis C virus (Meng, Yang et al. 2015). Consistently with the HIV-1 data, overexpression of the E2 ligases UBE2J2 and UBE2L6 promoted a significant inhibition of SINV_mCherry_ (fig. 6c). Overexpression of HNRNPD and POLDIP3, both containing IFN-α regulated phosphosites, also inhibited SINV gene expression (fig. 6c, suppl.f6f).

Most proteins caused a delay in SINV_mCherry_ that ranged from 1 to 2h, with EPSTI1, POLDIP3, and UBE2L6 promoting the strongest effects (fig. 6d). When extended to the 4∼20 hpi window, we noticed distinct delay patterns across candidates. EPSTI1 overexpression led to a continuous increase in Δt, whereas the delay with UBE2L6 overexpression remained constant (fig. 6e). We performed linear regression for Δt for the 12-20 hpi window and compared the different delay patterns based on the resulting slope (fig. 6f, top). Six candidates had clear positive slopes (NCOA4, USP41, THAP10, HNRNPD, POLDIP3, EPSTI1), suggesting that viral suppression increases over time (fig. 6f, bottom; suppl.f6i). Other proteins showed no substantial increase in Δt after initial inhibition. Altogether, these results imply two different behaviours: while some candidates exert their functions throughout the infection, others limit their action to the early stages of infection allowing viral gene expression recovery.

## DISCUSSION

In this study, we have analysed the proteome associated with different cellular states, including the permissive HEK293T, intermediate HEK293 and the hostile environment generated upon IFN-α stimulation. Our results highlight the widespread differences between HEK293T and HEK293 cells in their proteome, despite both lines having a shared lineage. Many laboratories have employed HEK293T cells due to their capacity to sustain high virus production, in analogy to other permissive cells such us Vero and BHK-21. Due to their high transfection efficiency and recombinant protein expression capacity, HEK293T cells have also been broadly employed to answer fundamental questions about the molecular biology of viruses (Liu, Sanchez et al. 2012, Schoggins, MacDuff et al. 2014, Hubel, Urban et al. 2019, Lei, Dong et al. 2020). Our results demonstrate that, in contrast to HEK293, the proteome of HEK293T lacks important cellular proteins that are central players in host-virus interactions, including a wide range of ISGs.

Conversely, the proteome changes in HEK293 associated with the establishment of the antiviral state by IFN-α are limited and focused exclusively on a group of core ISGs that are robustly expressed at the protein level. Most IFN-α studies have used microarrays and RNA-seq to survey the intracellular environment, revealing thousands of genes that respond to IFN-α (Rusinova, Forster et al. 2013). However, our results propose a more complex scenario in which a limited set of genes display a robust increase both at RNA and protein level, while many others show changes in RNA that are not matched at the protein level or *vice versa*. Interestingly, robustly expressed ISGs follow a defined pattern in which RNA expression is noticeable early upon IFN-α stimulation (here 4 hpt), while the transcriptional response consolidates into protein expression at later time points (here 20 hpt). Robustly expressed ISGs (both at mRNA and protein level) are strongly enriched in ISGs with proven capacity to supress infection (Schoggins, Wilson et al. 2011, Schoggins, MacDuff et al. 2014), and are conserved in the IFN response across mammalian species (Shaw, Hughes et al. 2017).

Conversely, a large group of genes displayed changes at the RNA level that are not reflected at the protein level. Discordance between protein and RNA have been observed in different cellular conditions (Beyer, Hollunder et al. 2004, Lee, Topper et al. 2011, Schwanhausser, Busse et al. 2011, Wilhelm, Schlegl et al. 2014, Jovanovic, Rooney et al. 2015), and our work demonstrates that it also occurs during the response to IFN-α. This must now be considered when interpreting experiments from IFN-treated cells using RNA centric approaches. Inconsistencies between RNA and protein can be explained by the diverse array of regulatory processed that collectively define gene expression, including RNA transcription, translation and turnover, as well as protein homeostasis (Liu, Beyer et al. 2016, Fortelny, Overall et al. 2017, Buccitelli and Selbach 2020). Conceptually, there are two possible biological explanations for protein/RNA discordance: i) transcriptional noise that has limited influence in the proteome because buffering mechanisms involving protein homeostasis that compensate for stochastic transcriptome fluctuations (McManus, May et al. 2014, Battle, Khan et al. 2015, Liu, Zhang et al. 2016, Eling, Morgan et al. 2019, Xiao, Hafner et al. 2021, Kusnadi, Timpone et al. 2022). This is likely the case of IFN-α downregulated genes as their overlapping across transcriptomic analyses is sparse, which is compatible with transcriptional noise (Shaw, Hughes et al. 2017, Colli, Ramos-Rodriguez et al. 2020). ii) The transcriptional induction of these genes occurs late upon IFN-α stimulation, possibly due to prolonged exposure. In this scenario, our experimental design would fail to capture derived proteome changes as it misses a later time point (e.g. ≥48 hpt). Notably, several of these late responder genes are involved in apoptosis, which suggests that production of the machinery to facilitate cell suicide is the last line of defence against virus infection.

Transcriptomic analysis of IFN-α also reveal a population of transcripts that are downregulated and have been associated with a higher CpG content (Shaw, Rihn et al. 2021). However, we did not observe an effect of the loss of these mRNAs at the proteome level at 20 hpt, suggesting the existence of mechanisms to maintain the homeostat of their encoded proteins. Conversely, 6 proteins had lower abundance after IFN-α treatment, while the levels of their cognate mRNAs remained unaltered. While these cases are small, our data suggests the existence of mechanisms that regulate protein stability, perhaps through ubiquitination pathways (Davis and Gack 2015).

The intracellular environment can also be regulated by subtle mechanisms that do not require changes in protein levels, such as the case for post-translational modifications. Our data reveal that as early as 10 min after interferon treatment, there are changes in the phosphorylation status of hundreds of phosphosites. The impact of rapid responses modulating the hostility of the intracellular environment is reflected in the data presented in suppl.f2a, where short IFN-α treatments cause substantial effects throughout the time course of SINV infection. Interestingly, many of these phosphorylation events involve RBPs, which are central players of virus infection, promoting and restricting infection (Garcia-Moreno, Jarvelin et al. 2018, Castello, Álvarez et al. 2024). Many of these phosphosites are placed near or within the RBDs of these RBPs. Given the negative charge of the phosphorylated amino acid and the highly negative nature of the phosphate backbone of RNA, these posttranslational modifications are expected to impair the ability of RBPs to bind RNA. However, it is difficult to distinguish between spurious and functionally relevant phosphosites, and further research should focus on characterising their functional impact individually.

Through integrated omics analysis, we identified a group of cellular proteins with no or poorly characterised antiviral roles. Our assays revealed virus-inhibitory effects for most proteins tested (13/15 for SINV and 6/9 for HIV-1), ranging from very mild to strong. It is worth noting that two candidate genes, HNRNPD and POLDIP3, were selected based on their IFN-α responses at the phosphorylation level. These two proteins have been reported to bind to the RNA of multiple types of viruses and have multifaceted roles in regulating virus replications (Paek, Kim et al. 2008, Lund, Milev et al. 2012, Hino, Sato et al. 2013, Liu, Yuan et al. 2017, Wu, Li et al. 2023). Our assays confirmed their inhibitory effects against SINV and HIV-1, but whether the IFN-α induced phosphorylation of these proteins is important in the infection phenotypes must be further investigated. Our results with SINV infection also revealed different inhibitory mechanisms, with some proteins targeting the early stages of virus infection (e.g. UBE2L6), and others participating throughout the course of infection (e.g. EPSTI1). The mechanism underlying these different kinetics of antiviral effects remains an interesting topic for future studies.

Our data provide a rich resource for studying determinants of virus permissiveness across cell states, and IFN-α response across transcriptome, proteome, and phosphoproteome. The proteome-wide differential expressions between HEK293 and more permissive HEK293T cell highlights the need to carefully distinguish these two cell lines in studies in virology, immunity and related fields. The multi-omic analysis shows that transcriptome variations cannot fully explain the observed proteome changes, calling for careful consideration and validation.

## ACKNOWLEDGEMENTS

A.C. is funded by the European Research Council (ERC) Consolidator Grant ‘vRNP-capture’ N# 101001634, the Career Development Award #MR/L019434/1, the John Fell Funds from the University of Oxford and the MRC grants MR/R021562/1 and MC_UU_00034/2. Q.G. and D.L.R. are funded by Medical Research Council grant MC_UU_00034/5. Antibody to HIV-1 p24 (ARP3279) was obtained from the Centre for AIDS Reagents, NIBSC, UK, supported by EURIPRED (EC FP7 INFRASTRUCTURES-2012 - INFRA-2012-1.1.5.: Grant Number 31266).

## DATA AVAILABILITY

The mass spectrometry proteomics data have been deposited to the ProteomeXchange Consortium via the PRIDE (Perez-Riverol, Bai et al. 2022) partner repository.

The RNA-seq data have been deposited to the GEO repository.

## MATERIALS AND METHODS

### Cell culture

The following human cell lines are available commercially available: HEK293 cell line (ECACC, #85120602), HEK293 Flp-In TRex cell line (Thermo Fisher Scientific, #R78007). HEK293T were kindly provided by Prof. Jan Rehwinkel (University of Oxford, UK). HEK293 Flp-In TRex inducible expression cells expressing proteins of interests were generated with Flp-In TRex Core Kit (Thermo Fisher Scientific, #K6500-01) according to manufacturer’s protocol, plasmids used for inducible cell line generation are described in following sections.

Cells were cultured in DMEM supplemented with 10 % fetal bovine serum (FBS) and 1x penicillin/streptomycin (Sigma Aldrich, #P4458) and the following specific antibiotics: 100 μg/ml Zeocin (Thermo Fisher Scientific, #R25001) and 7.5 μg/ml Blasticidin S Hydrochloride (Cambridge Bioscience, #B001-100mg) for HEK293 Flp-In TRex; 350 μg/ml Hygromycin B (Cambridge Bioscience, #H011-20ml) and 7.5 μg/ml Blasticidin for HEK293 Flp-In TRex inducible expression cells. All cells were cultured in a humidified incubator at 37 °C with 5% CO2. Culture with interferon alpha (IFN-a) stimulation was performed by supplying IFN-α stocks into culture media at indicated concentration and time period prior to harvest. IFN-α stocks were obtained by dissolving commercial IFN-α (PBL assay science, #11100-1) with 0.1% BSA in water and stored at −80°C.

### Cell culture in SILAC media

Cells were cultured in arginine and lysine depleted DMEM (Thermo Scientific, #10107883) supplied with 10% dialysed FBS (Silantes GmbH, #281000900), 1x penicillin/streptomycin, and isotope labeled arginine and lysine (Silantes GmbH amino acids: L-Arginine 13C,15N #201604102; L-Arginine 13C #201204102; L-Lysine 13C,15N #211604102; 4.4.5.5.-D4-L-Lysine #211104113). Cells were cultured in corresponding SILAC media for at least 5 more passages before experiments. Isotope incorporation rates over 99% were confirmed by mass spectrometry using whole cell lysates.

### Viruses

SINVmCherry suspension was generated with plasmid pT7-SVmCherry as described in our previous study (Garcia-Moreno, Noerenberg et al. 2019). Pseudotyped HIV-1Nef-mCherry and HIV-1Gag-mCherry were generated by co-transfection of HEK293T cells with pNL4-3. R-E-derived plasmids and a plasmid encoding the vesicular stomatitis virus glycoprotein (pHEF-VSVG, NIH AIDS Reagent Program, #4693) as described in our previous study (Garcia-Moreno, Noerenberg et al. 2019).

### Plasmids

Plasmids for producing inducible cell lines were generated by conventional cloning methods and gene synthesize: coding sequences of DTX3L and UBE2L6 were amplified from cDNA of IFN-a stimulated HEK293 using specific primers (key resources table), coding sequences of BTBD2, CMPK2, EPSTI1, FRMD5, HERC6, HNRNPD, LGALS3BP, MTRF1, NCOA4, POLDIP3, THAP10, UBE2J2, USP41 were generated by gene synthesis through Bio Basics Inc (suppl.tab5). Cloned and synthesized sequences were flanked with BamHI/HindIII/KpnI at 5’ and NotI at 3’ ends restriction sites and cloned into the vector pcDNA5/FRT/TO with eGFP preceded or followed by a Gly-Ser linker sequence (GGSGGSGG).

### mCherry-based virus fitness assay

Cells were seeded on 96-well microplate (flat-bottom, Greiner Bio-One, #655986) at 6x104 cells per well, in complete DMEM (lack phenol-red) supplied with 5% FBS, 1 mM sodium pyruvate, 1x penicillin/streptomycin. For IFN-a stimulation assays, IFN-a was supplemented to each well at specified concentration at least 4 hours after cell seeding. For Tet-ON cells, 1 μg/ml doxycycline was supplied in culture media during cell seeding. Infection of SINVmCherry, HIV-1Gag-mCherry or HIV-1Nef-mCherry was performed at least 20 hours after cell seeding with complete DMEM (lack phenol-red) containing 2.5% FBS, 1 mM sodium pyruvate, 1x penicillin/streptomycin, and virus at specified MOI. Cells were incubated at 37 °C and 5% CO2 in a CLARIOstar fluorescence plate reader (BMG Labtech) for 24 h (SINV) or 48h (HIV-1). Fluorescence results were obtained with n ≥ 9 for SINV assays (3 replicates per plate, ≥ 3 plates) and n ≥ 6 for HIV-1 assays (3 replicates per plate, ≥ 2 plates).

Fluorescence signals collected from CLARIOstar plate reader were analysed with R. For analysis of statistical significance, mCherry signals were scaled to 0-100 to account for between-plate variations, then determined by t test. For analysis of fluorescence signal delay with mCherry-tagged viruses, the average of scaled mCherry signals at each timepoint was calculated for each plate, and the fluorescence signal delay (Δt) at each timepoint on each plate was obtained by finding the earliest timepoint that the average signal in overexpressed cells exceed that in control cells, these Δt values were then summarised into Δt-over-hpi plots by plotting the average and standard deviation of at each Δt timepoint. Analysis of slopes of Δt-over-hpi curve was performed with linear regression using R.

### Proteomics sample preparation

SILAC samples for HEK293 versus HEK293T cells proteomics analysis were cultured on 6-well plate (2 light + 1 heavy samples for HEK293, 2 heavy + 1 light samples for HEK293T); samples for HEK293 mock versus IFN-a stimulated cells proteomics and phosphoproteomics analysis were cultured on 150 cm2 dishes (1 light + 1 medium + 1 heavy samples for mock, IFN-a 10min, IFN-a 4h, and IFN-a 20h). Cells were lysed on plate with urea lysis buffer (8 M urea (Sigma, U1250), 100 mM ammonium bicarbonate (AmBic; Sigma, #9830), 10 μl/ml protease inhibitor cocktail (Sigma, P8340) and 2.5 μl/ml phosphatase inhibitor (Sigma, P0044) after gentle washes with cold PBS. Protein concentration in each sample was measured with Pierce 660-nm Protein Assay (Thermo Scientific, #22660) followed by sample mixing with label swap.

For proteomics analysis, combined lysates were reduced and alkylated with addition of 10 nM Tris(2-carboxyethyl)phosphine hydrochloride (TCEP; Thermo Scientific #77720) and 50 mM 2-Chloroacetamide (CAA; Sigma, #C0267) and in-dark incubation for 30 min. Alkylated samples were diluted to 6 M urea with 25 mM AmBic for LysC protease (Wako, #129-02541) digestion with protein/protease ratio at 40:1 (w/w) at 37 °C for 4 h, followed by dilution to 1 M urea with 25 mM AmBic and 37 °C o/n digestion with trypsin protease (MS grade; Promega, V5280) in protein/protease ratio of 40:1 (w/w). Digestion was quenched with 0.5% (v/v) formic acid (FA; Thermo Scientific, A117-50) and frozen in −80 °C prior to fractionation. Tryptic peptide fractionation was performed with off-line HPLC, loaded with solvent A (10 mM AmBic 2% acetonitrile (ACN; Sigma, #34851) in water, pH 8.3) and separated by a Zorbax 300 Extended-C18 column (2.1 x 150 mm, 3.5 μm) using a linear gradient (length: 100 min, 8% to 60% solvent B (80% ACN in water), flow rate: 200 μl/min). Fractions were collected every 1-min interval between 12 - 92 min of the gradient. The 80 collected fractions were combined at an even interval (1st, 21st, 41st, 61st fractions were combined, and so on), resulting in 20 fractions. Fractions were dried with SpeedVac and stored in −80 °C, and reconstituted with loading buffer (5% DMSO, 5% FA in water) prior to LC-MSMS analysis.

For phosphoproteomics analysis, combined samples were reduced and alkylated with addition of 10 mM TCEP and 50 mM CAA and in-dark incubation for 30 min. Alkylated samples were purified with methanol-chloroform precipitation by sequential supply of methanol, chloroform, water in 4:1:3 ratio (v/v) with vortex after each reagent addition, followed by centrifugation in 3,000 g for 10 min and removal of all liquid. Remaining samples were air dried and reconstituted in 8 M urea buffer. Purified alkylated proteins were diluted to 6 M urea with 25 mM AmBic for LysC protease digestion with protein/protease ratio at 60:1 (w/w) at 37 °C for 4 h, followed by dilution to 1 M urea with 25 mM AmBic and 37 °C o/n digestion with trypsin protease in protein/protease ratio of 50:1 (w/w), then quenched with 0.5% (v/v) FA. Fractionation of tryptic peptide was performed with the same method as proteomics samples resulting in 20 fractions, which were dried with Speed Vac and resuspended in 125 μl loading buffer, then combined into 10 fractions at an even interval (1st with 11th, 2nd with 12th, etc.). These 10 fractions were subjected to phosphopeptide enrichment by Ti-IMAC (Resyn Bioscience, MR-TIM002) according to manufacturer’s protocol. Final eluates were dried with SpeedVac and stored in in −80 °C prior to LC-MSMS analysis.

### Mass spectrometry

For proteomics analysis, reconstituted tryptic peptides were analysed on a nanoUHPLC (Thermo) connected to a Q Exactive mass spectrometer (Thermo Fischer Scientific) through an EASY-Spray nano-electrospray ion source (Thermo Fischer Scientific). The peptides were trapped on a C18 PepMap100 pre-column (300 µm i.d. x 5 mm, 100 Å, Thermo Fisher) using solvent A (0.1% formic acid in water). The peptides were separated on an EASY-spray Acclaim PepMap analytical column (75 µm i.d. × 500 mm, RSLC C18, 2 µm, 100 Å) using a linear gradient (length: 120 min, 8 % to 28 % solvent B (0.1% formic acid, 5 % DMSO in acetonitrile), flow rate: 200 nL/min). The separated peptides were electrosprayed directly into the mass spectrometer operating in a data-dependent mode. Full scan MS spectra were acquired in the Orbitrap (scan range 350-1500 m/z, resolution 70000, AGC target 3e6, maximum injection time 100 ms). After the MS scans, the 20 most intense peaks were selected for HCD fragmentation at 30 % of normalised collision energy. HCD spectra were also acquired in the Orbitrap (resolution 17500, AGC target 5e4, maximum injection time 120 ms).

For phosphoproteomics analysis, Ti-IMAC enriched phosphopeptides were reconstituted in loading buffer and analysed on two instruments. One half of reconstituted phosphopeptides were analysed on a nanoUHPLC (Thermo) connected to a Q Exactive mass spectrometer (Thermo Fischer Scientific) through an EASY-Spray nano-electrospray ion source (Thermo Fischer Scientific). The peptides were trapped on a C18 PepMap100 pre-column (300 µm i.d. x 5 mm, 100 Å, Thermo Fisher) using solvent A (0.1% formic acid in water). The peptides were separated on an EASY-spray Acclaim PepMap analytical column (75 µm i.d. × 500 mm, RSLC C18, 2 µm, 100 Å) using a linear gradient (length: 120 min, 8 % to 28 % solvent B (0.1% formic acid, 5 % DMSO in acetonitrile), flow rate: 200 nL/min). The separated peptides were electrosprayed directly into the mass spectrometer operating in a data-dependent mode. Full scan MS spectra were acquired in the Orbitrap (scan range 350-1500 m/z, resolution 70000, AGC target 3e6, maximum injection time 50 ms). After the MS scans, the 20 most intense peaks were selected for HCD fragmentation at 30 % of normalised collision energy. HCD spectra were also acquired in the Orbitrap (resolution 17500, AGC target 5e4, maximum injection time 120 ms). The other half of phosphopeptide samples were analysed on an EASY-nLC 1000 System (Thermo) connected to an Orbitrap Elite Hybrid mass spectrometer (Thermo Fischer Scientific) through an EASY-Spray nano-electrospray ion source (Thermo Fischer Scientific). The peptides were trapped on a C18 PepMap100 pre-column (300 µm i.d. x 5 mm, 100 Å, Thermo Fisher) using solvent A (0.1% formic acid in water). The peptides were separated on an EASY-spray Acclaim PepMap analytical column (75 µm i.d. × 500 mm, RSLC C18, 2 µm, 100 Å) using a linear gradient (length: 60 min, 8 % to 28 % solvent B (0.1% formic acid, 5 % DMSO in acetonitrile), flow rate: 200 nL/min). The separated peptides were electrosprayed directly into the mass spectrometer operating in a data-dependent mode. Full scan MS spectra were acquired in the Orbitrap (scan range 350-1500 m/z, resolution 70000, AGC target 1e6, maximum injection time 100 ms). After the MS scans, the 20 most intense peaks were selected for CID fragmentation at 35% normalised collision energy. CID spectra were acquired in the LTQ mass analyser (resolution 7500, AGC target 5e3, maximum injection time 100 ms).

Protein identification and quantification were performed using Andromeda search engine implemented in MaxQuant (1.6.3.4). Peptides were searched Human Uniprot database (Uniprot_id: UP000005640, downloaded Nov 2016). Multiplicity was set to 2 for HEK293-versus-HEK293T proteomics analysis, specifying Arg10/Lys8 for heavy labels; while multiplicity was set to 3 for HEK293-versus-IFN-stimulated HEK293 proteomics and phosphoproteomics analysis, specifying Arg6/Lys4 for medium labels and Arg10/Lys8 for heavy labels. False discovery rate (FDR) was set at 1% for both peptide and protein identification. Phospho(STY) was set as a variable modification for phosphoproteomics analysis, and raw files from two instruments were searched separately. Data files from the same sample were assigned as near-by fractions, and match between run was turned on to match peak information from near-by fractions. Other options were set as defaults.

### Proteomics quantitative analysis

For protein quantification, proteinGroups files from MaxQuant search outputs were used for quantitative analysis. Analysis was performed in R (v.4.3.2, R Core Team). Proteins flagged as decoy or potential contaminant were filtered out. Proteins with at least 3 valid values in at least one condition were subjected to log 2 transformation, normalisation, and imputation of missing values. Normalisation was performed with vsn method using package vsn (Huber, von Heydebreck et al. 2002), imputation was performed by background-level signal using deterministic minimum method (minDet) in package ImputeLCMD (Lazar 2015). Principal component analysis (PCA) was performed to evaluate batch effects prior to imputation. Statistical analysis was performed using moderated t test in package limma (Ritchie, Phipson et al. 2015). For HEK293-HEK293T proteomics analysis, replicate number was set as block variable (Smyth, Michaud et al. 2005); for HEK293 mock-IFN-a proteomics analysis, replicate number was set as block variable and isotope type was modelled as co-variable in design matrix. P values were adjusted for multiple-testing with Benjamini-Hochberg method.

For phosphosite quantification, Phospho(STY) files from MaxQuant search outputs were used for quantitative analysis. Phosphosites with at least 3 valid values in at least one condition were subjected to log 2 transformation, normalisation, and imputation of missing values. Normalisation was performed with vsn method using package vsn, imputation was performed by background-level signal using deterministic minimum method (minDet) in package ImputeLCMD. Principal component analysis (PCA) was performed to evaluate batch effects prior to imputation. Statistical analysis included both moderated t test and Z test. Moderated t test was performed using package limma with replicate number set as block variable and isotope type modelled as co-variable. Z test was performed by obtaining Z-score via dividing average fold change of each phosphosite by the global standard deviation, then convert Z-scores to p-values. All p values were adjusted for multiple-testing with Benjamini-Hochberg method. For phosphosites that did not meet quantification criteria, sites detected with 2 valid values in one condition and completely missing in the other condition were selected for semi-quantitative analysis by calculating intensity, fold change, and rank among total semi-quantitation sites.

Phosphoproteome data from two instruments was tested separately and these analysis results are available in suppl.tab3 “Separate result table”. Test results from two instruments were further combined following these rules: 1), all sites only quantified in one instrument were accepted; 2), for sites quantified in both instruments, accept the result with higher localisation probability; 3), if the difference between localisation probabilities was less than 0.05, accept the result with higher identification score; 4), if both results scored over 90, accept the result with higher quantile in the dataset. The combined phosphosite quantification result is available in suppl.tab3 “Combined result table”.

### RNA sequencing

HEK293 cells were cultured on 6-well plates in triplicate with IFN-a treatments (mock, IFN-a 4h, IFN-a 20h). Total RNA was extracted from cells using Monarch Total RNA Miniprep Kit (NEB, T2010S) following the manufacturer’s protocol. Briefly, cells were gently washed with 4 °C PBS and lysed with 600 μl RNA Lysis Buffer provided in the RNA miniprep kit with gentle pipetting. Genomic DNA (gDNA) were removed using gDNA Removal Column with centrifuge, flow-throughs were supplied with ethanol and loaded on RNA Extract Columns with centrifuge. RNA on column was washed with Washing Buffer, then eluted with 100 μl nuclease-free water. Ribosomal RNAs were removed by rRNA Depletion Kit (NEB, E6310) following the manufacturer’s protocol. Briefly, total RNA was hybridised with rRNA probes in rRNA Depletion Solution, then sequentially digested by RNase H and DNase I at 37 °C for 30 min each. Extracted RNAs were supplied with RiboLock RNase inhibitor (Thermo Scientific, EO0381) and subjected to SuperScript III reverse transcriptase (Invitrogen, #18080093) for cDNA library generation using random hexamer primers. RNase H was added to reverse transcription products to obtain cDNA library for RNA-seq. Libraries were pooled in equimolar concentrations and sequenced using a NextSeq 500 sequencer (Illumina).

RNA-Seq reads quality was assessed using FastQC software (http://www.bioinformatics.babraham.ac.uk/projects/fastqc). 89.1% of the reads generated presented a Q score of ≥30. RNA-Seq reads were aligned to the Homo sapiens genome (GRCh38) downloaded via ENSEMBL using Hisat2 (Kim, Langmead et al. 2015). After the alignment, software FeatureCount was used to count reads mapping to genes annotation files (Liao, Smyth et al. 2014). For differential expression analysis of RNA-seq results, the edgeR package was used to measure gene expression levels by normalizing the raw counts to counts per million (CPM) and to identify differentially expressed genes between sample groups (Robinson, McCarthy et al. 2010).

### Bioinformatic analysis

Gene set enrichment analysis (GSEA) and pathway enrichment were performed with STRING online platform (string-db.org) (Szklarczyk, Gable et al. 2019). Further analysis was performed in R (v.4.3.2, R Core Team). Enrichments of viral protein and viral RNA interactors in proteomics results were obtained by first matching significant proteins to a manually compiled compilation of viral protein/RNA interactors, then calculate enrichment with Fisher’s exact test. Viral protein interactors were compiled based on series of published large-scale GFP-pulldown experiments (Jager, Cimermancic et al. 2012, Pichlmair, Kandasamy et al. 2012, Shah, Link et al. 2018, Gordon, Jang et al. 2020). Viral RNA interactors were obtained from our recent review article (Iselin, Palmalux et al. 2022). Enrichment analysis of viral protein/RNA interactors in phosphoproteomics results was performed by first obtaining proteins that contain IFN-a regulated phosphosites (FDR<0.05 in moderted T or Z statistics), then match to the compilation of viral protein/RNA interactors described above and calculated with Fisher’s exact test.

Matching of IFN-related GO annotation was performed with package GO.db (Carlson 2020) and biomaRt (Steffen Durinck 2009). GO terms containing keywords “innate immune / type I interferon / interferon-alpha” were considered IFN-related terms. Genes annotated with these terms were extracted and matched to proteomics results. Matching of ISG interactors was performed using proteins with FDR<0.05 in large-scale proteomic-based interactor screening study (Hubel, Urban et al. 2019). Matching of FACS-screened ISGs was performed using proteins with Z < −1.5 from screening results (Schoggins, Wilson et al. 2011, Schoggins, MacDuff et al. 2014).

Kinase-substrate enrichment analysis (KSEA) was performed following the procedure described in published studies (Lachmann and Ma’ayan 2009, Bouhaddou, Memon et al. 2020). Briefly, kinase-substrate correlation annotations were manually compiled from series of online databases (PhosphoSitePlus (Hornbeck, Kornhauser et al. 2012), PhosphoELM (Dinkel, Chica et al. 2011), SIGNOR (Licata, Lo Surdo et al. 2020), DEPOD (Damle and Kohn 2019), iPTMnet (Huang, Arighi et al. 2018), NetworKIN and Netphorest (Linding, Jensen et al. 2008)). After matching to phosphoproteome quantification and semi-quantification results, the activity of each kinase was calculated by the average fold change of phosphosites that it can regulate. Kinases of significantly altered activities were determined by Z-statistics of all kinase activity values, with cutoff set at FDR < 0.05.

Distances between phosphosites and RNA binding domains (RBDs) were calculated based on RNA-interactive peptide fragments (RBDpep) in RBDmap (Castello, Fischer et al. 2016). Briefly, RBDpep peptide fragments in each RBP were assembled into non-overlapping RNA binding regions, the distance of a site was set zero if it locates inside RNA binding regions, and determined as the number of amino acid between site and closest RNA binding region. Distances of random distributed sites were calculated as follows: for sites locate inside RNA binding region, distances were remained zero; for sites locate between two RNA binding regions, distances were calculated as expected values when site positions distribute randomly within those sequences.

## Key resources table

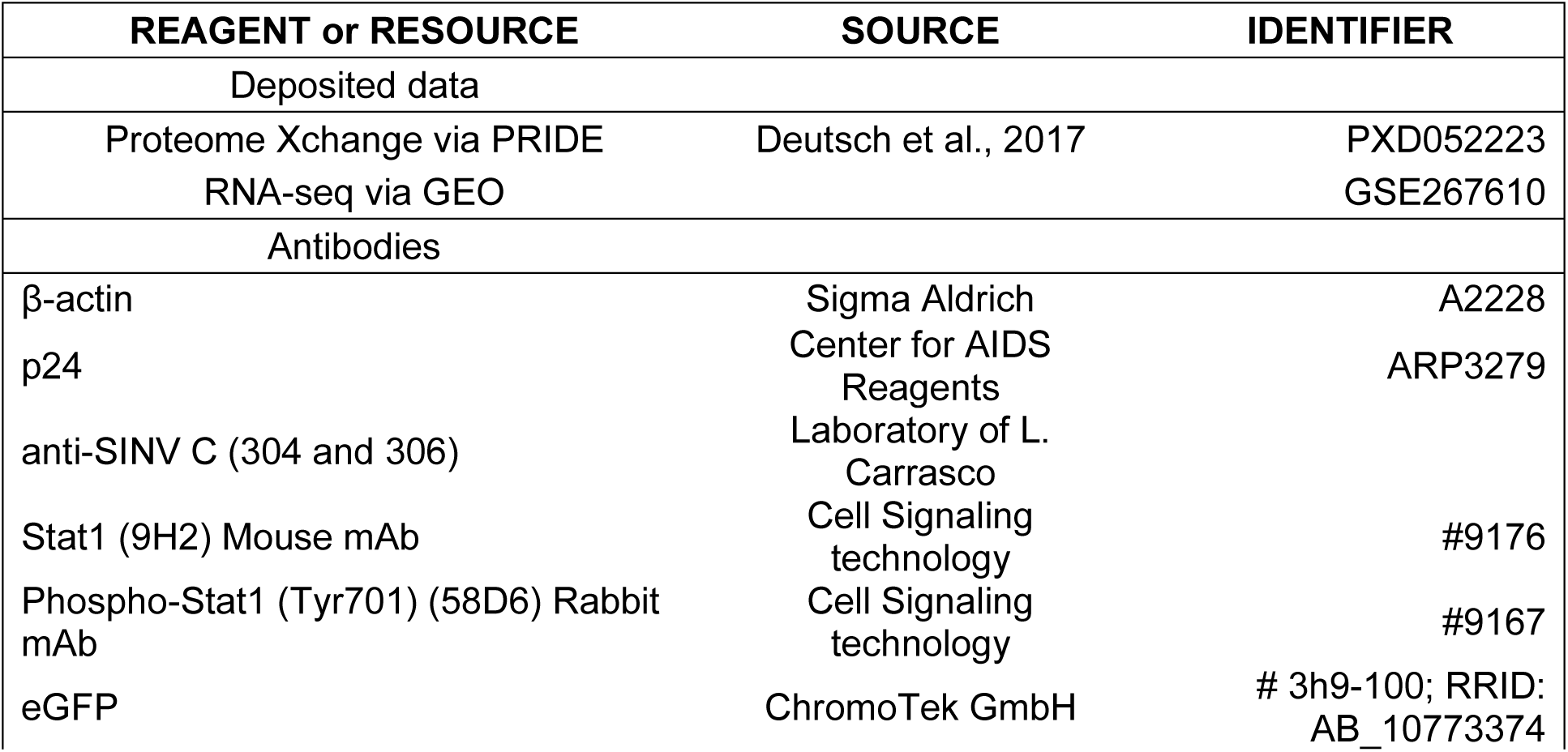
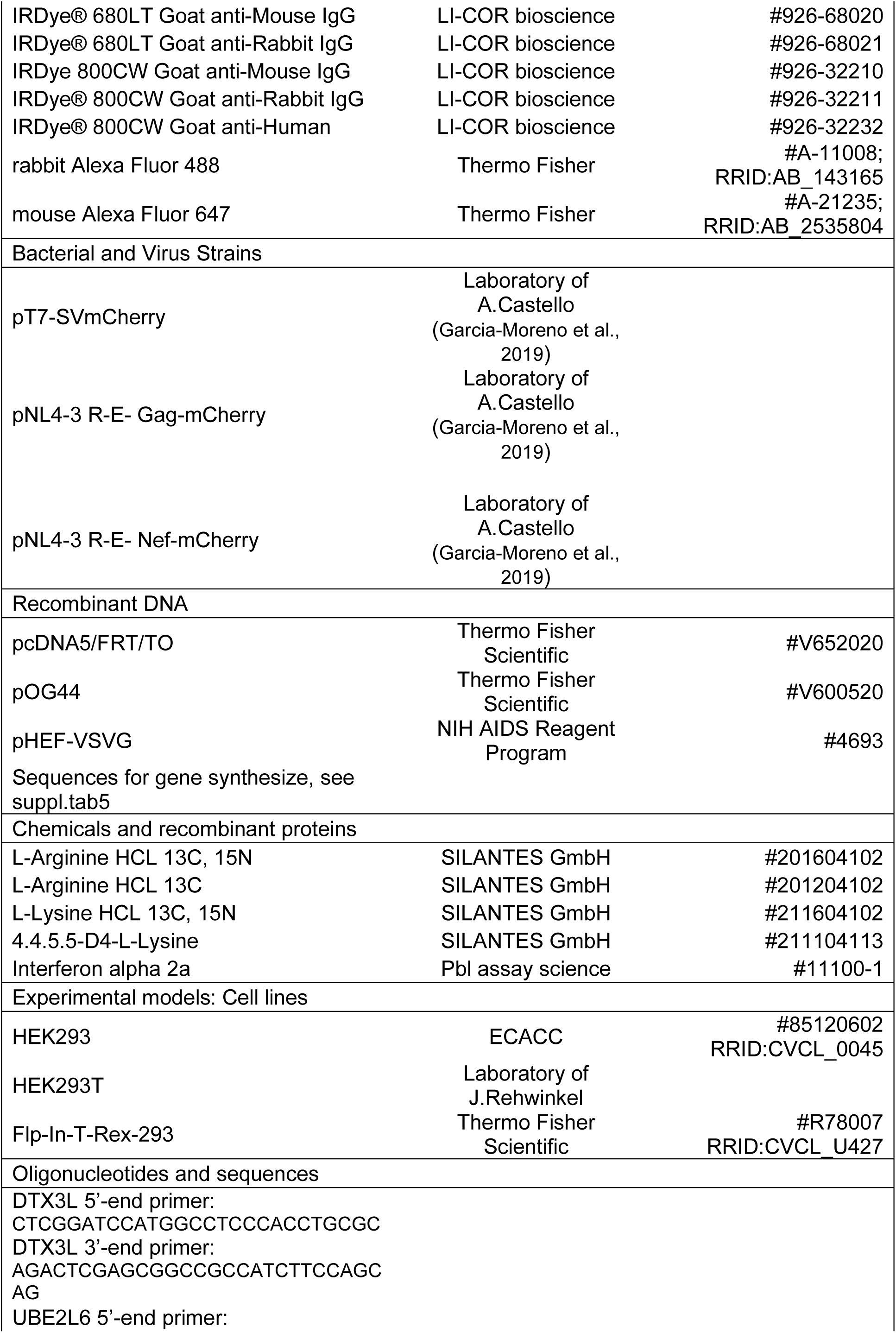
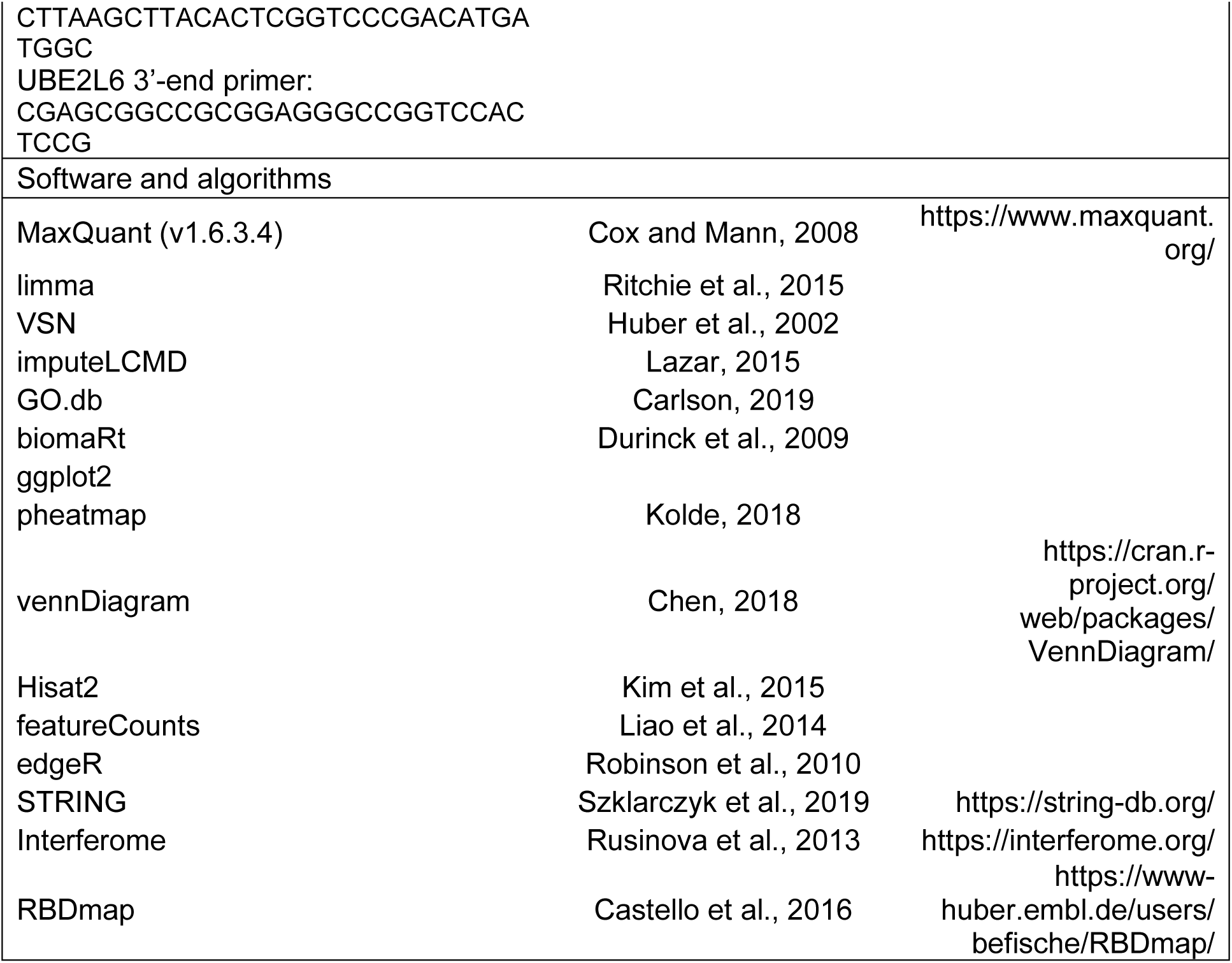

## REFERENCES

Arhel, N., A. Genovesio, K. A. Kim, S. Miko, E. Perret, J. C. Olivo-Marin, S. Shorte and P. Charneau (2006). “Quantitative four-dimensional tracking of cytoplasmic and nuclear HIV-1 complexes.” Nat Methods 3(10): 817–824.

Bae, D. H., M. Marino, B. Iaffaldano, S. Fenstermaker, S. Afione, T. Argaw, J. McCright, A. Kwilas, J. A. Chiorini, A. E. Timmons and J. Reiser (2020). “Design and Testing of Vector-Producing HEK293T Cells Bearing a Genomic Deletion of the SV40 T Antigen Coding Region.” Molecular Therapy-Methods & Clinical Development 18: 631–638.

Barbalat, R., S. E. Ewald, M. L. Mouchess and G. M. Barton (2011). “Nucleic acid recognition by the innate immune system.” Annu Rev Immunol 29: 185–214.

Battle, A., Z. Khan, S. H. Wang, A. Mitrano, M. J. Ford, J. K. Pritchard and Y. Gilad (2015). “Genomic variation. Impact of regulatory variation from RNA to protein.” Science 347(6222): 664–667.

Beyer, A., J. Hollunder, H. P. Nasheuer and T. Wilhelm (2004). “Post-transcriptional expression regulation in the yeast Saccharomyces cerevisiae on a genomic scale.” Mol Cell Proteomics 3(11): 1083–1092.

Blees, A., D. Januliene, T. Hofmann, N. Koller, C. Schmidt, S. Trowitzsch, A. Moeller and R. Tampe (2017). “Structure of the human MHC-I peptide-loading complex.” Nature 551(7681): 525–528.

Boone, M., Y. C. Lin, L. Meuris, I. Lemmens, N. Van Roy, A. Soete, J. Reumers, M. Moisse, S. Plaisance, R. Drmanac, J. Chen, F. Speleman, D. Lambrechts, Y. Van de Peer, J. Tavernier and N. Callewaert (2014). “Genome dynamics of the human embryonic kidney 293 (HEK293) lineage in response to cell biology manipulations.” New Biotechnology 31: S71–S72.

Bouhaddou, M., D. Memon, B. Meyer, K. M. White, V. V. Rezelj, M. Correa Marrero, B. J. Polacco, J. E. Melnyk, S. Ulferts, R. M. Kaake, J. Batra, A. L. Richards, E. Stevenson, D. E. Gordon, A. Rojc, K. Obernier, J. M. Fabius, M. Soucheray, L. Miorin, E. Moreno, C. Koh, Q. D. Tran, A. Hardy, R. Robinot, T. Vallet, B. E. Nilsson-Payant, C. Hernandez-Armenta, A. Dunham, S. Weigang, J. Knerr, M. Modak, D. Quintero, Y. Zhou, A. Dugourd, A. Valdeolivas, T. Patil, Q. Li, R. Huttenhain, M. Cakir, M. Muralidharan, M. Kim, G. Jang, B. Tutuncuoglu, J. Hiatt, J. Z. Guo, J. Xu, S. Bouhaddou, C. J. P. Mathy, A. Gaulton, E. J. Manners, E. Felix, Y. Shi, M. Goff, J. K. Lim, T. McBride, M. C. O’Neal, Y. Cai, J. C. J. Chang, D. J. Broadhurst, S. Klippsten, E. De Wit, A. R. Leach, T. Kortemme, B. Shoichet, M. Ott, J. Saez-Rodriguez, B. R. tenOever, R. D. Mullins, E. R. Fischer, G. Kochs, R. Grosse, A. Garcia-Sastre, M. Vignuzzi, J. R. Johnson, K. M. Shokat, D. L. Swaney, P. Beltrao and N. J. Krogan (2020). “The Global Phosphorylation Landscape of SARS-CoV-2 Infection.” Cell 182(3): 685–712 e619.

Brass, A. L., D. M. Dykxhoorn, Y. Benita, N. Yan, A. Engelman, R. J. Xavier, J. Lieberman and S. J. Elledge (2008). “Identification of host proteins required for HIV infection through a functional genomic screen.” Science 319(5865): 921–926.

Buccitelli, C. and M. Selbach (2020). “mRNAs, proteins and the emerging principles of gene expression control.” Nat Rev Genet 21(10): 630–644.

Burgui, I., T. Aragon, J. Ortin and A. Nieto (2003). “PABP1 and eIF4GI associate with influenza virus NS1 protein in viral mRNA translation initiation complexes.” J Gen Virol 84(Pt 12): 3263–3274.

Bushell, M. and P. Sarnow (2002). “Hijacking the translation apparatus by RNA viruses.” J Cell Biol 158(3): 395–399.

Busse, D. C., D. Habgood-Coote, S. Clare, C. Brandt, I. Bassano, M. Kaforou, J. Herberg, M. Levin, J. F. Eleouet, P. Kellam and J. S. Tregoning (2020). “Interferon-Induced Protein 44 and Interferon-Induced Protein 44-Like Restrict Replication of Respiratory Syncytial Virus.” J Virol 94(18).

Carlson, M. (2020). GO.db: A set of annotation maps describing the entire Gene Ontology.

Castello, A., L. Álvarez, W. Kamel, L. Iselin and J. Hennig (2024). “Exploring the Expanding Universe of Host-Virus Interactions Mediated by Viral RNA.” Mol Cell.

Castello, A., B. Fischer, C. K. Frese, R. Horos, A. M. Alleaume, S. Foehr, T. Curk, J. Krijgsveld and M. W. Hentze (2016). “Comprehensive Identification of RNA-Binding Domains in Human Cells.” Mol Cell 63(4): 696–710.

Chen, J., K. Sathiyamoorthy, X. Zhang, S. Schaller, B. E. Perez White, T. S. Jardetzky and R. Longnecker (2018). “Ephrin receptor A2 is a functional entry receptor for Epstein-Barr virus.” Nat Microbiol 3(2): 172–180.

Cheng, K., L. Martin-Sancho, L. R. Pal, Y. Pu, L. Riva, X. Yin, S. Sinha, N. U. Nair, S. K. Chanda and E. Ruppin (2021). “Genome-scale metabolic modeling reveals SARS-CoV-2-induced metabolic changes and antiviral targets.” Mol Syst Biol 17(11): e10260.

Colli, M. L., M. Ramos-Rodriguez, E. S. Nakayasu, M. I. Alvelos, M. Lopes, J. L. E. Hill, J. V. Turatsinze, A. Coomans de Brachene, M. A. Russell, H. Raurell-Vila, A. Castela, J. Juan-Mateu, B. M. Webb-Robertson, L. Krogvold, K. Dahl-Jorgensen, L. Marselli, P. Marchetti, S. J. Richardson, N. G. Morgan, T. O. Metz, L. Pasquali and D. L. Eizirik (2020). “An integrated multi-omics approach identifies the landscape of interferon-alpha-mediated responses of human pancreatic beta cells.” Nat Commun 11(1): 2584.

Damle, N. P. and M. Kohn (2019). “The human DEPhOsphorylation Database DEPOD: 2019 update.” Database (Oxford) 2019.

Darnell, J. E., Jr. (1997). “STATs and gene regulation.” Science 277(5332): 1630–1635.

Davis, M. E. and M. U. Gack (2015). “Ubiquitination in the antiviral immune response.” Virology 479-480: 52–65.

De, B. P., S. Cram, H. Lee, J. B. Rosenberg, D. Sondhi, R. G. Crystal and S. M. Kaminsky (2023). “Assessment of Residual Full-Length SV40 Large T Antigen in Clinical-Grade Adeno-Associated Virus Vectors Produced in 293T Cells.” Hum Gene Ther 34(15-16): 697–704.

Di Palma, S., M. L. Hennrich, A. J. Heck and S. Mohammed (2012). “Recent advances in peptide separation by multidimensional liquid chromatography for proteome analysis.” J Proteomics 75(13): 3791–3813.

Dinkel, H., C. Chica, A. Via, C. M. Gould, L. J. Jensen, T. J. Gibson and F. Diella (2011). “Phospho.ELM: a database of phosphorylation sites--update 2011.” Nucleic Acids Res 39(Database issue): D261–267.

Donovan, J., M. Dufner and A. Korennykh (2013). “Structural basis for cytosolic double-stranded RNA surveillance by human oligoadenylate synthetase 1.” Proc Natl Acad Sci U S A 110(5): 1652–1657.

Dragic, T., V. Litwin, G. P. Allaway, S. R. Martin, Y. Huang, K. A. Nagashima, C. Cayanan, P. J. Maddon, R. A. Koup, J. P. Moore and W. A. Paxton (1996). “HIV-1 entry into CD4+ cells is mediated by the chemokine receptor CC-CKR-5.” Nature 381(6584): 667–673.

Dunne, A., M. Ejdeback, P. L. Ludidi, L. A. O’Neill and N. J. Gay (2003). “Structural complementarity of Toll/interleukin-1 receptor domains in Toll-like receptors and the adaptors Mal and MyD88.” J Biol Chem 278(42): 41443–41451.

El Bougrini, J., L. Dianoux and M. K. Chelbi-Alix (2011). “PML positively regulates interferon gamma signaling.” Biochimie 93(3): 389–398.

Eling, N., M. D. Morgan and J. C. Marioni (2019). “Challenges in measuring and understanding biological noise.” Nat Rev Genet 20(9): 536–548.

Fan, T. J., L. H. Han, R. S. Cong and J. Liang (2005). “Caspase family proteases and apoptosis.” Acta Biochim Biophys Sin (Shanghai) 37(11): 719–727.

Ferreira, C. B., R. P. Sumner, M. T. Rodriguez-Plata, J. Rasaiyaah, R. S. Milne, A. J. Thrasher, W. Qasim and G. J. Towers (2020). “Lentiviral vector production titer is not limited in HEK293T by induced intracellular innate immunity.” Molecular Therapy-Methods & Clinical Development 17: 209–219.

Floyd-Smith, G., E. Slattery and P. Lengyel (1981). “Interferon action: RNA cleavage pattern of a (2’-5’)oligoadenylate--dependent endonuclease.” Science 212(4498): 1030–1032.

Forero, A., N. S. Giacobbi, K. D. McCormick, O. V. Gjoerup, C. J. Bakkenist, J. M. Pipas and S. N. Sarkar (2014). “Simian Virus 40 Large T Antigen Induces IFN-Stimulated Genes through ATR Kinase.” Journal of Immunology 192(12): 5933–5942.

Fortelny, N., C. M. Overall, P. Pavlidis and G. V. C. Freue (2017). “Can we predict protein from mRNA levels?” Nature 547(7664): E19–E20.

Gack, M. U., Y. C. Shin, C. H. Joo, T. Urano, C. Liang, L. Sun, O. Takeuchi, S. Akira, Z. Chen, S. Inoue and J. U. Jung (2007). “TRIM25 RING-finger E3 ubiquitin ligase is essential for RIG-I-mediated antiviral activity.” Nature 446(7138): 916–920.

Garcia-Moreno, M., A. I. Jarvelin and A. Castello (2018). “Unconventional RNA-binding proteins step into the virus-host battlefront.” Wiley Interdiscip Rev RNA 9(6): e1498.

Garcia-Moreno, M., M. Noerenberg, S. Ni, A. I. Jarvelin, E. Gonzalez-Almela, C. E. Lenz, M. Bach-Pages, V. Cox, R. Avolio, T. Davis, S. Hester, T. J. M. Sohier, B. Li, G. Heikel, G. Michlewski, M. A. Sanz, L. Carrasco, E. P. Ricci, V. Pelechano, I. Davis, B. Fischer, S. Mohammed and A. Castello (2019). “System-wide Profiling of RNA-Binding Proteins Uncovers Key Regulators of Virus Infection.” Mol Cell 74(1): 196–211 e111.

Garcia-Moreno, M., R. Truman, H. Chen, L. Iselin, C. E. Lenz, J. Y. Lee, K. Dicker, M. Noerenberg, T. J. M. Sohier, N. Palmalux, A. I. Jarvelin, W. Kamel, V. Ruscica, E. P. Ricci, I. Davis, S. Mohammed and A. Castello (2023). “Incorporation of genome-bound cellular proteins into HIV-1 particles regulates viral infection.” BioRxiv.

Garcia, M. A., E. F. Meurs and M. Esteban (2007). “The dsRNA protein kinase PKR: virus and cell control.” Biochimie 89(6-7): 799–811.

Gordon, D. E., G. M. Jang, M. Bouhaddou, J. Xu, K. Obernier, K. M. White, M. J. O’Meara, V. V. Rezelj, J. Z. Guo, D. L. Swaney, T. A. Tummino, R. Huttenhain, R. M. Kaake, A. L. Richards, B. Tutuncuoglu, H. Foussard, J. Batra, K. Haas, M. Modak, M. Kim, P. Haas, B. J. Polacco, H. Braberg, J. M. Fabius, M. Eckhardt, M. Soucheray, M. J. Bennett, M. Cakir, M. J. McGregor, Q. Li, B. Meyer, F. Roesch, T. Vallet, A. Mac Kain, L. Miorin, E. Moreno, Z. Z. C. Naing, Y. Zhou, S. Peng, Y. Shi, Z. Zhang, W. Shen, I. T. Kirby, J. E. Melnyk, J. S. Chorba, K. Lou, S. A. Dai, I. Barrio-Hernandez, D. Memon, C. Hernandez-Armenta, J. Lyu, C. J. P. Mathy, T. Perica, K. B. Pilla, S. J. Ganesan, D. J. Saltzberg, R. Rakesh, X. Liu, S. B. Rosenthal, L. Calviello, S. Venkataramanan, J. Liboy-Lugo, Y. Lin, X. P. Huang, Y. Liu, S. A. Wankowicz, M. Bohn, M. Safari, F. S. Ugur, C. Koh, N. S. Savar, Q. D. Tran, D. Shengjuler, S. J. Fletcher, M. C. O’Neal, Y. Cai, J. C. J. Chang, D. J. Broadhurst, S. Klippsten, P. P. Sharp, N. A. Wenzell, D. Kuzuoglu-Ozturk, H. Y. Wang, R. Trenker, J. M. Young, D. A. Cavero, J. Hiatt, T. L. Roth, U. Rathore, A. Subramanian, J. Noack, M. Hubert, R. M. Stroud, A. D. Frankel, O. S. Rosenberg, K. A. Verba, D. A. Agard, M. Ott, M. Emerman, N. Jura, M. von Zastrow, E. Verdin, A. Ashworth, O. Schwartz, C. d’Enfert, S. Mukherjee, M. Jacobson, H. S. Malik, D. G. Fujimori, T. Ideker, C. S. Craik, S. N. Floor, J. S. Fraser, J. D. Gross, A. Sali, B. L. Roth, D. Ruggero, J. Taunton, T. Kortemme, P. Beltrao, M. Vignuzzi, A. Garcia-Sastre, K. M. Shokat, B. K. Shoichet and N. J. Krogan (2020). “A SARS-CoV-2 protein interaction map reveals targets for drug repurposing.” Nature 583(7816): 459–468.

Gu, L., A. Fullam, R. Brennan and M. Schroder (2013). “Human DEAD box helicase 3 couples IkappaB kinase epsilon to interferon regulatory factor 3 activation.” Mol Cell Biol 33(10): 2004–2015.

Harada, H., T. Fujita, M. Miyamoto, Y. Kimura, M. Maruyama, A. Furia, T. Miyata and T. Taniguchi (1989). “Structurally similar but functionally distinct factors, IRF-1 and IRF-2, bind to the same regulatory elements of IFN and IFN-inducible genes.” Cell 58(4): 729–739.

Harcourt, J., A. Tamin, X. Lu, S. Kamili, S. K. Sakthivel, J. Murray, K. Queen, Y. Tao, C. R. Paden, J. Zhang, Y. Li, A. Uehara, H. Wang, C. Goldsmith, H. A. Bullock, L. Wang, B. Whitaker, B. Lynch, R. Gautam, C. Schindewolf, K. G. Lokugamage, D. Scharton, J. A. Plante, D. Mirchandani, S. G. Widen, K. Narayanan, S. Makino, T. G. Ksiazek, K. S. Plante, S. C. Weaver, S. Lindstrom, S. Tong, V. D. Menachery and N. J. Thornburg (2020). “Severe Acute Respiratory Syndrome Coronavirus 2 from Patient with Coronavirus Disease, United States.” Emerg Infect Dis 26(6): 1266–1273.

Henig, N., N. Avidan, I. Mandel, E. Staun-Ram, E. Ginzburg, T. Paperna, R. Y. Pinter and A. Miller (2013). “Interferon-beta induces distinct gene expression response patterns in human monocytes versus T cells.” PLoS One 8(4): e62366.

Hino, K., H. Sato, A. Sugai, M. Kato, M. Yoneda and C. Kai (2013). “Downregulation of Nipah virus N mRNA occurs through interaction between its 3’ untranslated region and hnRNP D.” J Virol 87(12): 6582–6588.

Hornbeck, P. V., J. M. Kornhauser, S. Tkachev, B. Zhang, E. Skrzypek, B. Murray, V. Latham and M. Sullivan (2012). “PhosphoSitePlus: a comprehensive resource for investigating the structure and function of experimentally determined post-translational modifications in man and mouse.” Nucleic Acids Res 40(Database issue): D261–270.

Hou, F., L. Sun, H. Zheng, B. Skaug, Q. X. Jiang and Z. J. Chen (2011). “MAVS forms functional prion-like aggregates to activate and propagate antiviral innate immune response.” Cell 146(3): 448–461.

Huang, H., C. N. Arighi, K. E. Ross, J. Ren, G. Li, S. C. Chen, Q. Wang, J. Cowart, K. Vijay-Shanker and C. H. Wu (2018). “iPTMnet: an integrated resource for protein post-translational modification network discovery.” Nucleic Acids Res 46(D1): D542–D550.

Hubel, P., C. Urban, V. Bergant, W. M. Schneider, B. Knauer, A. Stukalov, P. Scaturro, A. Mann, L. Brunotte, H. H. Hoffmann, J. W. Schoggins, M. Schwemmle, M. Mann, C. M. Rice and A. Pichlmair (2019). “A protein-interaction network of interferon-stimulated genes extends the innate immune system landscape.” Nat Immunol 20(4): 493–502.

Huber, W., A. von Heydebreck, H. Sultmann, A. Poustka and M. Vingron (2002). “Variance stabilization applied to microarray data calibration and to the quantification of differential expression.” Bioinformatics 18 **Suppl 1**: S96–104.

Iselin, L., N. Palmalux, W. Kamel, P. Simmonds, S. Mohammed and A. Castello (2022). “Uncovering viral RNA-host cell interactions on a proteome-wide scale.” Trends in Biochemical Sciences 47(1): 23–38.

Jager, S., P. Cimermancic, N. Gulbahce, J. R. Johnson, K. E. McGovern, S. C. Clarke, M. Shales, G. Mercenne, L. Pache, K. Li, H. Hernandez, G. M. Jang, S. L. Roth, E. Akiva, J. Marlett, M. Stephens, I. D’Orso, J. Fernandes, M. Fahey, C. Mahon, A. J. O’Donoghue, A. Todorovic, J. H. Morris, D. A. Maltby, T. Alber, G. Cagney, F. D. Bushman, J. A. Young, S. K. Chanda, W. I. Sundquist, T. Kortemme, R. D. Hernandez, C. S. Craik, A. Burlingame, A. Sali, A. D. Frankel and N. J. Krogan (2012). “Global landscape of HIV-human protein complexes.” Nature 481(7381): 365–370.

Jovanovic, M., M. S. Rooney, P. Mertins, D. Przybylski, N. Chevrier, R. Satija, E. H. Rodriguez, A. P. Fields, S. Schwartz, R. Raychowdhury, M. R. Mumbach, T. Eisenhaure, M. Rabani, D. Gennert, D. Lu, T. Delorey, J. S. Weissman, S. A. Carr, N. Hacohen and A. Regev (2015). “Immunogenetics. Dynamic profiling of the protein life cycle in response to pathogens.” Science 347(6226): 1259038.

Kato, H., O. Takeuchi, S. Sato, M. Yoneyama, M. Yamamoto, K. Matsui, S. Uematsu, A. Jung, T. Kawai, K. J. Ishii, O. Yamaguchi, K. Otsu, T. Tsujimura, C. S. Koh, C. Reis e Sousa, Y. Matsuura, T. Fujita and S. Akira (2006). “Differential roles of MDA5 and RIG-I helicases in the recognition of RNA viruses.” Nature 441(7089): 101–105.

Kim, B., S. Arcos, K. Rothamel, J. Jian, K. L. Rose, W. H. McDonald, Y. Bian, S. Reasoner, N. J. Barrows, S. Bradrick, M. A. Garcia-Blanco and M. Ascano (2020). “Discovery of Widespread Host Protein Interactions with the Pre-replicated Genome of CHIKV Using VIR-CLASP.” Mol Cell 78(4): 624–640 e627.

Kim, D., B. Langmead and S. L. Salzberg (2015). “HISAT: a fast spliced aligner with low memory requirements.” Nat Methods 12(4): 357–360.

Kusnadi, E. P., C. Timpone, I. Topisirovic, O. Larsson and L. Furic (2022). “Regulation of gene expression via translational buffering.” Biochim Biophys Acta Mol Cell Res 1869(1): 119140.

Lachmann, A. and A. Ma’ayan (2009). “KEA: kinase enrichment analysis.” Bioinformatics 25(5): 684–686.

Lai, J. H., D. W. Wu, C. H. Wu, L. F. Hung, C. Y. Huang, S. M. Ka, A. Chen, Z. F. Chang and L. J. Ho (2021). “Mitochondrial CMPK2 mediates immunomodulatory and antiviral activities through IFN-dependent and IFN-independent pathways.” iScience 24(6): 102498.

Lau, L., E. E. Gray, R. L. Brunette and D. B. Stetson (2015). “DNA tumor virus oncogenes antagonize the cGAS-STING DNA-sensing pathway.” Science 350(6260): 568–571.

Lazar, C. (2015). “imputeLCMD: a collection of methods for left-censored missing data imputation.” R package, version 2.

Lee, M. V., S. E. Topper, S. L. Hubler, J. Hose, C. D. Wenger, J. J. Coon and A. P. Gasch (2011). “A dynamic model of proteome changes reveals new roles for transcript alteration in yeast.” Mol Syst Biol 7: 514.

Lei, X., X. Dong, R. Ma, W. Wang, X. Xiao, Z. Tian, C. Wang, Y. Wang, L. Li, L. Ren, F. Guo, Z. Zhao, Z. Zhou, Z. Xiang and J. Wang (2020). “Activation and evasion of type I interferon responses by SARS-CoV-2.” Nat Commun 11(1): 3810.

Li, B., S. M. Clohisey, B. S. Chia, B. Wang, A. Cui, T. Eisenhaure, L. D. Schweitzer, P. Hoover, N. J. Parkinson, A. Nachshon, N. Smith, T. Regan, D. Farr, M. U. Gutmann, S. I. Bukhari, A. Law, M. Sangesland, I. Gat-Viks, P. Digard, S. Vasudevan, D. Lingwood, D. H. Dockrell, J. G. Doench, J. K. Baillie and N. Hacohen (2020). “Genome-wide CRISPR screen identifies host dependency factors for influenza A virus infection.” Nat Commun 11(1): 164.

Li, M. Q., E. Kao, X. Gao, H. Sandig, K. Limmer, M. Pavon-Eternod, T. E. Jones, S. Landry, T. Pan, M. D. Weitzman and M. David (2012). “Codon-usage-based inhibition of HIV protein synthesis by human schlafen 11.” Nature 491(7422): 125–U145.

Li, Q., A. L. Brass, A. Ng, Z. Hu, R. J. Xavier, T. J. Liang and S. J. Elledge (2009). “A genome-wide genetic screen for host factors required for hepatitis C virus propagation.” Proc Natl Acad Sci U S A 106(38): 16410–16415.

Li, Y., R. Chen, Q. Zhou, Z. Xu, C. Li, S. Wang, A. Mao, X. Zhang, W. He and H. B. Shu (2012). “LSm14A is a processing body-associated sensor of viral nucleic acids that initiates cellular antiviral response in the early phase of viral infection.” Proc Natl Acad Sci U S A 109(29): 11770–11775.

Liao, Y., G. K. Smyth and W. Shi (2014). “featureCounts: an efficient general purpose program for assigning sequence reads to genomic features.” Bioinformatics 30(7): 923–930.

Licata, L., P. Lo Surdo, M. Iannuccelli, A. Palma, E. Micarelli, L. Perfetto, D. Peluso, A. Calderone, L. Castagnoli and G. Cesareni (2020). “SIGNOR 2.0, the SIGnaling Network Open Resource 2.0: 2019 update.” Nucleic Acids Res 48(D1): D504–D510.

Lin, Y. C., M. Boone, L. Meuris, I. Lemmens, N. Van Roy, A. Soete, J. Reumers, M. Moisse, S. Plaisance, R. Drmanac, J. Chen, F. Speleman, D. Lambrechts, Y. Van de Peer, J. Tavernier and N. Callewaert (2014). “Genome dynamics of the human embryonic kidney 293 lineage in response to cell biology manipulations.” Nature Communications 5.

Linding, R., L. J. Jensen, A. Pasculescu, M. Olhovsky, K. Colwill, P. Bork, M. B. Yaffe and T. Pawson (2008). “NetworKIN: a resource for exploring cellular phosphorylation networks.” Nucleic Acids Res 36(Database issue): D695–699.

Liu, S. Y., D. J. Sanchez, R. Aliyari, S. Lu and G. Cheng (2012). “Systematic identification of type I and type II interferon-induced antiviral factors.” Proc Natl Acad Sci U S A 109(11): 4239–4244.

Liu, T., J. Zhang and T. Zhou (2016). “Effect of Interaction between Chromatin Loops on Cell-to-Cell Variability in Gene Expression.” PLoS Comput Biol 12(5): e1004917.

Liu, X. N., J. H. Yuan, T. T. Wang, W. Pan and S. H. Sun (2017). “An alternative POLDIP3 transcript promotes hepatocellular carcinoma progression.” Biomed Pharmacother 89: 276–283.

Liu, Y., A. Beyer and R. Aebersold (2016). “On the Dependency of Cellular Protein Levels on mRNA Abundance.” Cell 165(3): 535–550.

Lund, N., M. P. Milev, R. Wong, T. Sanmuganantham, K. Woolaway, B. Chabot, S. Abou Elela, A. J. Mouland and A. Cochrane (2012). “Differential effects of hnRNP D/AUF1 isoforms on HIV-1 gene expression.” Nucleic Acids Res 40(8): 3663–3675.

McManus, C. J., G. E. May, P. Spealman and A. Shteyman (2014). “Ribosome profiling reveals post-transcriptional buffering of divergent gene expression in yeast.” Genome Res 24(3): 422–430.

Meng, X., D. Yang, R. Yu and H. Zhu (2015). “EPSTI1 Is Involved in IL-28A-Mediated Inhibition of HCV Infection.” Mediators Inflamm 2015: 716315.

Merten, O. W., S. Charrier, N. Laroudie, S. Fauchille, C. Dugue, C. Jenny, M. Audit, M. A. Zanta-Boussif, H. Chautard, M. Radrizzani, G. Vallanti, L. Naldini, P. Noguiez-Hellin and A. Galy (2011). “Large-Scale Manufacture and Characterization of a Lentiviral Vector Produced for Clinical Ex Vivo Gene Therapy Application.” Human Gene Therapy 22(3): 343–356.

Meyer, K., Y. C. Kwon, S. Liu, C. H. Hagedorn, R. B. Ray and R. Ray (2015). “Interferon-alpha inducible protein 6 impairs EGFR activation by CD81 and inhibits hepatitis C virus infection.” Sci Rep 5: 9012.

Milian, E., T. Julien, R. Biaggio, A. Venereo-Sanchez, J. Montes, A. P. Manceur, S. Ansorge, E. Petiot, M. Rosa-Calatrava and A. Kamen (2017). “Accelerated mass production of influenza virus seed stocks in HEK-293 suspension cell cultures by reverse genetics.” Vaccine 35(26): 3423–3430.

Modrof, J., A. Kerschbaum, M. R. Farcet, D. Niemeyer, V. M. Corman and T. R. Kreil (2020). “SARS-CoV-2 and the safety margins of cell-based biological medicinal products.” Biologicals 68: 122–124.

Needham, E. J., B. L. Parker, T. Burykin, D. E. James and S. J. Humphrey (2019). “Illuminating the dark phosphoproteome.” Sci Signal 12(565).

Ong, S. E., B. Blagoev, I. Kratchmarova, D. B. Kristensen, H. Steen, A. Pandey and M. Mann (2002). “Stable isotope labeling by amino acids in cell culture, SILAC, as a simple and accurate approach to expression proteomics.” Mol Cell Proteomics 1(5): 376–386.

Paek, K. Y., C. S. Kim, S. M. Park, J. H. Kim and S. K. Jang (2008). “RNA-binding protein hnRNP D modulates internal ribosome entry site-dependent translation of hepatitis C virus RNA.” J Virol 82(24): 12082–12093.

Park, J. Y., B. P. Lim, K. Lee, Y. G. Kim and E. C. Jo (2006). “Scalable production of adeno-associated virus type 2 vectors via suspension transfection.” Biotechnol Bioeng 94(3): 416–430.

Perez-Riverol, Y., J. Bai, C. Bandla, D. Garcia-Seisdedos, S. Hewapathirana, S. Kamatchinathan, D. J. Kundu, A. Prakash, A. Frericks-Zipper, M. Eisenacher, M. Walzer, S. Wang, A. Brazma and J. A. Vizcaino (2022). “The PRIDE database resources in 2022: a hub for mass spectrometry-based proteomics evidences.” Nucleic Acids Res 50(D1): D543–D552.

Pichlmair, A., K. Kandasamy, G. Alvisi, O. Mulhern, R. Sacco, M. Habjan, M. Binder, A. Stefanovic, C. A. Eberle, A. Goncalves, T. Burckstummer, A. C. Muller, A. Fauster, C. Holze, K. Lindsten, S. Goodbourn, G. Kochs, F. Weber, R. Bartenschlager, A. G. Bowie, K. L. Bennett, J. Colinge and G. Superti-Furga (2012). “Viral immune modulators perturb the human molecular network by common and unique strategies.” Nature 487(7408): 486–U101.

Pichlmair, A., O. Schulz, C. P. Tan, T. I. Naslund, P. Liljestrom, F. Weber and C. Reis e Sousa (2006). “RIG-I-mediated antiviral responses to single-stranded RNA bearing 5’-phosphates.” Science 314(5801): 997–1001.

Platanias, L. C. (2005). “Mechanisms of type-I- and type-II-interferon-mediated signalling.” Nat Rev Immunol 5(5): 375–386.

Pritchard, L. K., D. J. Harvey, C. Bonomelli, M. Crispin and K. J. Doores (2015). “Cell- and Protein-Directed Glycosylation of Native Cleaved HIV-1 Envelope.” J Virol 89(17): 8932–8944.

Radoshitzky, S. R., G. Pegoraro, X. O. Chi, D. N. L, C. Y. Chiang, L. Jozwick, J. C. Clester, C. L. Cooper, D. Courier, D. P. Langan, K. Underwood, K. A. Kuehl, M. G. Sun, Y. Cai, S. Q. Yu, R. Burk, R. Zamani, K. Kota, J. H. Kuhn and S. Bavari (2016). “siRNA Screen Identifies Trafficking Host Factors that Modulate Alphavirus Infection.” PLoS Pathog 12(3): e1005466.

Reus, J. B., G. S. Trivino-Soto, L. I. Wu, K. Kokott and E. S. Lim (2020). “SV40 large T antigen is not responsible for the loss of STING in 293T cells but can inhibit cGAS-STING interferon induction.” Viruses 12(2): 137.

Rio, D. C., S. G. Clark and R. Tjian (1985). “A mammalian host-vector system that regulates expression and amplification of transfected genes by temperature induction.” Science 227(4682): 23–28.

Ritchie, M. E., B. Phipson, D. Wu, Y. F. Hu, C. W. Law, W. Shi and G. K. Smyth (2015). “limma powers differential expression analyses for RNA-sequencing and microarray studies.” Nucleic Acids Research 43(7).

Ritter, J. B., A. S. Wahl, S. Freund, Y. Genzel and U. Reichl (2010). “Metabolic effects of influenza virus infection in cultured animal cells: Intra- and extracellular metabolite profiling.” BMC Syst Biol 4: 61.

Robinson, M. D., D. J. McCarthy and G. K. Smyth (2010). “edgeR: a Bioconductor package for differential expression analysis of digital gene expression data.” Bioinformatics 26(1): 139–140.

Rusinova, I., S. Forster, S. Yu, A. Kannan, M. Masse, H. Cumming, R. Chapman and P. J. Hertzog (2013). “INTERFEROME v2.0: an updated database of annotated interferon-regulated genes.” Nucleic Acids Research 41(D1): D1040–D1046.

Sadler, A. J. and B. R. Williams (2008). “Interferon-inducible antiviral effectors.” Nat Rev Immunol 8(7): 559–568.

Savidis, G., W. M. McDougall, P. Meraner, J. M. Perreira, J. M. Portmann, G. Trincucci, S. P. John, A. M. Aker, N. Renzette, D. R. Robbins, Z. Guo, S. Green, T. F. Kowalik and A. L. Brass (2016). “Identification of Zika Virus and Dengue Virus Dependency Factors using Functional Genomics.” Cell Rep 16(1): 232–246.

Schmidt, N., S. Ganskih, Y. Wei, A. Gabel, S. Zielinski, H. Keshishian, C. A. Lareau, L. Zimmermann, J. Makroczyova, C. Pearce, K. Krey, T. Hennig, S. Stegmaier, L. Moyon, M. Horlacher, S. Werner, J. Aydin, M. Olguin-Nava, R. Potabattula, A. Kibe, L. Dolken, R. P. Smyth, N. Caliskan, A. Marsico, C. Krempl, J. Bodem, A. Pichlmair, S. A. Carr, P. Chlanda, F. Erhard and M. Munschauer (2023). “SND1 binds SARS-CoV-2 negative-sense RNA and promotes viral RNA synthesis through NSP9.” Cell 186(22): 4834–4850 e4823.

Schneider, W. M., M. D. Chevillotte and C. M. Rice (2014). “Interferon-stimulated genes: a complex web of host defenses.” Annu Rev Immunol 32: 513–545.

Schoggins, J. W., D. A. MacDuff, N. Imanaka, M. D. Gainey, B. Shrestha, J. L. Eitson, K. B. Mar, R. B. Richardson, A. V. Ratushny, V. Litvak, R. Dabelic, B. Manicassamy, J. D. Aitchison, A. Aderem, R. M. Elliott, A. Garcia-Sastre, V. Racaniello, E. J. Snijder, W. M. Yokoyama, M. S. Diamond, H. W. Virgin and C. M. Rice (2014). “Pan-viral specificity of IFN-induced genes reveals new roles for cGAS in innate immunity.” Nature 505(7485): 691–695.

Schoggins, J. W., S. J. Wilson, M. Panis, M. Y. Murphy, C. T. Jones, P. Bieniasz and C. M. Rice (2011). “A diverse range of gene products are effectors of the type I interferon antiviral response.” Nature 472(7344): 481–485.

Schwanhausser, B., D. Busse, N. Li, G. Dittmar, J. Schuchhardt, J. Wolf, W. Chen and M. Selbach (2011). “Global quantification of mammalian gene expression control.” Nature 473(7347): 337–342.

Shah, P. S., N. Link, G. M. Jang, P. P. Sharp, T. T. Zhu, D. L. Swaney, J. R. Johnson, J. Von Dollen, H. R. Ramage, L. Satkamp, B. Newton, R. Huttenhain, M. J. Petit, T. Baum, A. Everitt, O. Laufman, M. Tassetto, M. Shales, E. Stevenson, G. N. Iglesias, L. Shokat, S. Tripathi, V. Balasubramaniam, L. G. Webb, S. Aguirre, A. J. Willsey, A. Garcia-Sastre, K. S. Pollard, S. Cherry, A. V. Gamarnik, I. Marazzi, J. Taunton, A. Fernandez-Sesma, H. J. Bellen, R. Andino and N. J. Krogan (2018). “Comparative Flavivirus-Host Protein Interaction Mapping Reveals Mechanisms of Dengue and Zika Virus Pathogenesis.” Cell 175(7): 1931-+.

Shaw, A. E., J. Hughes, Q. Gu, A. Behdenna, J. B. Singer, T. Dennis, R. J. Orton, M. Varela, R. J. Gifford, S. J. Wilson and M. Palmarini (2017). “Fundamental properties of the mammalian innate immune system revealed by multispecies comparison of type I interferon responses.” PLoS Biol 15(12): e2004086.

Shaw, A. E., S. J. Rihn, N. Mollentze, A. Wickenhagen, D. G. Stewart, R. J. Orton, S. Kuchi, S. Bakshi, M. R. Collados, M. L. Turnbull, J. Busby, Q. Gu, K. Smollett, C. G. G. Bamford, E. Sugrue, P. C. D. Johnson, A. F. Da Silva, A. Castello, D. G. Streicker, D. L. Robertson, M. Palmarini and S. J. Wilson (2021). “The antiviral state has shaped the CpG composition of the vertebrate interferome to avoid self-targeting.” PLoS Biol 19(9): e3001352.

Silvennoinen, O., J. N. Ihle, J. Schlessinger and D. E. Levy (1993). “Interferon-induced nuclear signalling by Jak protein tyrosine kinases.” Nature 366(6455): 583–585.

Silverman, R. H. (2007). “Viral encounters with 2’,5’-oligoadenylate synthetase and RNase L during the interferon antiviral response.” J Virol 81(23): 12720–12729.

Smyth, G. K., J. Michaud and H. S. Scott (2005). “Use of within-array replicate spots for assessing differential expression in microarray experiments.” Bioinformatics 21(9): 2067–2075.

Soonsawad, P., L. Xing, E. Milla, J. M. Espinoza, M. Kawano, M. Marko, C. Hsieh, H. Furukawa, M. Kawasaki, W. Weerachatyanukul, R. Srivastava, S. W. Barnett, I. K. Srivastava and R. H. Cheng (2010). “Structural evidence of glycoprotein assembly in cellular membrane compartments prior to Alphavirus budding.” J Virol 84(21): 11145–11151.

Steffen Durinck, P. T. S., Ewan Birney and Wolfgang Huber (2009). “Mapping identifiers for the integration of genomic datasets with the R/Bioconductor package biomaRt.” Nature Protocols 4: 7.

Stremlau, M., C. M. Owens, M. J. Perron, M. Kiessling, P. Autissier and J. Sodroski (2004). “The cytoplasmic body component TRIM5alpha restricts HIV-1 infection in Old World monkeys.” Nature 427(6977): 848–853.

Sun, L., J. Wu, F. Du, X. Chen and Z. J. Chen (2013). “Cyclic GMP-AMP synthase is a cytosolic DNA sensor that activates the type I interferon pathway.” Science 339(6121): 786–791.

Sundquist, W. I. and H. G. Krausslich (2012). “HIV-1 assembly, budding, and maturation.” Cold Spring Harb Perspect Med 2(7): a006924.

Swaminathan, S., P. Rajan, O. Savinova, R. Jagus and B. Thimmapaya (1996). “Simian virus 40 large-T bypasses the translational block imposed by the phosphorylation of eIF-2 alpha.” Virology 219(1): 321–323.

Szklarczyk, D., A. L. Gable, D. Lyon, A. Junge, S. Wyder, J. Huerta-Cepas, M. Simonovic, N. T. Doncheva, J. H. Morris, P. Bork, L. J. Jensen and C. V. Mering (2019). “STRING v11: protein-protein association networks with increased coverage, supporting functional discovery in genome-wide experimental datasets.” Nucleic Acids Res 47(D1): D607–D613.

Tanaka, T., M. Hosokawa, V. V. Vagin, M. Reuter, E. Hayashi, A. L. Mochizuki, K. Kitamura, H. Yamanaka, G. Kondoh, K. Okawa, S. Kuramochi-Miyagawa, T. Nakano, R. Sachidanandam, G. J. Hannon, R. S. Pillai, N. Nakatsuji and S. Chuma (2011). “Tudor domain containing 7 (Tdrd7) is essential for dynamic ribonucleoprotein (RNP) remodeling of chromatoid bodies during spermatogenesis.” Proc Natl Acad Sci U S A 108(26): 10579–10584.

Wang, K. S., R. J. Kuhn, E. G. Strauss, S. Ou and J. H. Strauss (1992). “High-affinity laminin receptor is a receptor for Sindbis virus in mammalian cells.” J Virol 66(8): 4992–5001.

Wang, Y., X. Tong, G. Li, J. Li, M. Deng and X. Ye (2012). “Ankrd17 positively regulates RIG-I-like receptor (RLR)-mediated immune signaling.” Eur J Immunol 42(5): 1304–1315.

Wilhelm, M., J. Schlegl, H. Hahne, A. M. Gholami, M. Lieberenz, M. M. Savitski, E. Ziegler, L. Butzmann, S. Gessulat, H. Marx, T. Mathieson, S. Lemeer, K. Schnatbaum, U. Reimer, H. Wenschuh, M. Mollenhauer, J. Slotta-Huspenina, J. H. Boese, M. Bantscheff, A. Gerstmair, F. Faerber and B. Kuster (2014). “Mass-spectrometry-based draft of the human proteome.” Nature 509(7502): 582–587.

Winkler, R., E. Gillis, L. Lasman, M. Safra, S. Geula, C. Soyris, A. Nachshon, J. Tai-Schmiedel, N. Friedman, V. T. K. Le-Trilling, M. Trilling, M. Mandelboim, J. H. Hanna, S. Schwartz and N. Stern-Ginossar (2019). “m(6)A modification controls the innate immune response to infection by targeting type I interferons.” Nat Immunol 20(2): 173–182.

Wong, J. J., Y. F. Pung, N. S. Sze and K. C. Chin (2006). “HERC5 is an IFN-induced HECT-type E3 protein ligase that mediates type I IFN-induced ISGylation of protein targets.” Proc Natl Acad Sci U S A 103(28): 10735–10740.

Wu, Y., M. Li, J. Tian, H. Yan, Y. Pan, H. Shi, D. Shi, J. Chen, L. Guo and L. Feng (2023). “Broad antagonism of coronaviruses nsp5 to evade the host antiviral responses by cleaving POLDIP3.” PLoS Pathog 19(10): e1011702.

Xiao, J. Y., A. Hafner and A. N. Boettiger (2021). “How subtle changes in 3D structure can create large changes in transcription.” Elife 10.

Xu, G., Z. Xia, F. Deng, L. Liu, Q. Wang, Y. Yu, F. Wang, C. Zhu, W. Liu, Z. Cheng, Y. Zhu, L. Zhou, Y. Zhang, M. Lu and S. Liu (2019). “Inducible LGALS3BP/90K activates antiviral innate immune responses by targeting TRAF6 and TRAF3 complex.” PLoS Pathog 15(8): e1008002.

Xue, B., D. Yang, J. Wang, Y. Xu, X. Wang, Y. Qin, R. Tian, S. Chen, Q. Xie, N. Liu and H. Zhu (2016). “ISG12a Restricts Hepatitis C Virus Infection through the Ubiquitination-Dependent Degradation Pathway.” J Virol 90(15): 6832–6845.

Zhao, C., S. L. Beaudenon, M. L. Kelley, M. B. Waddell, W. Yuan, B. A. Schulman, J. M. Huibregtse and R. M. Krug (2004). “The UbcH8 ubiquitin E2 enzyme is also the E2 enzyme for ISG15, an IFN-alpha/beta-induced ubiquitin-like protein.” Proc Natl Acad Sci U S A 101(20): 7578–7582.

