## Supplementary figures for "Multi-omics analysis of virus-permissive versus hostile cellular states reveals protein networks controlling virus infection"

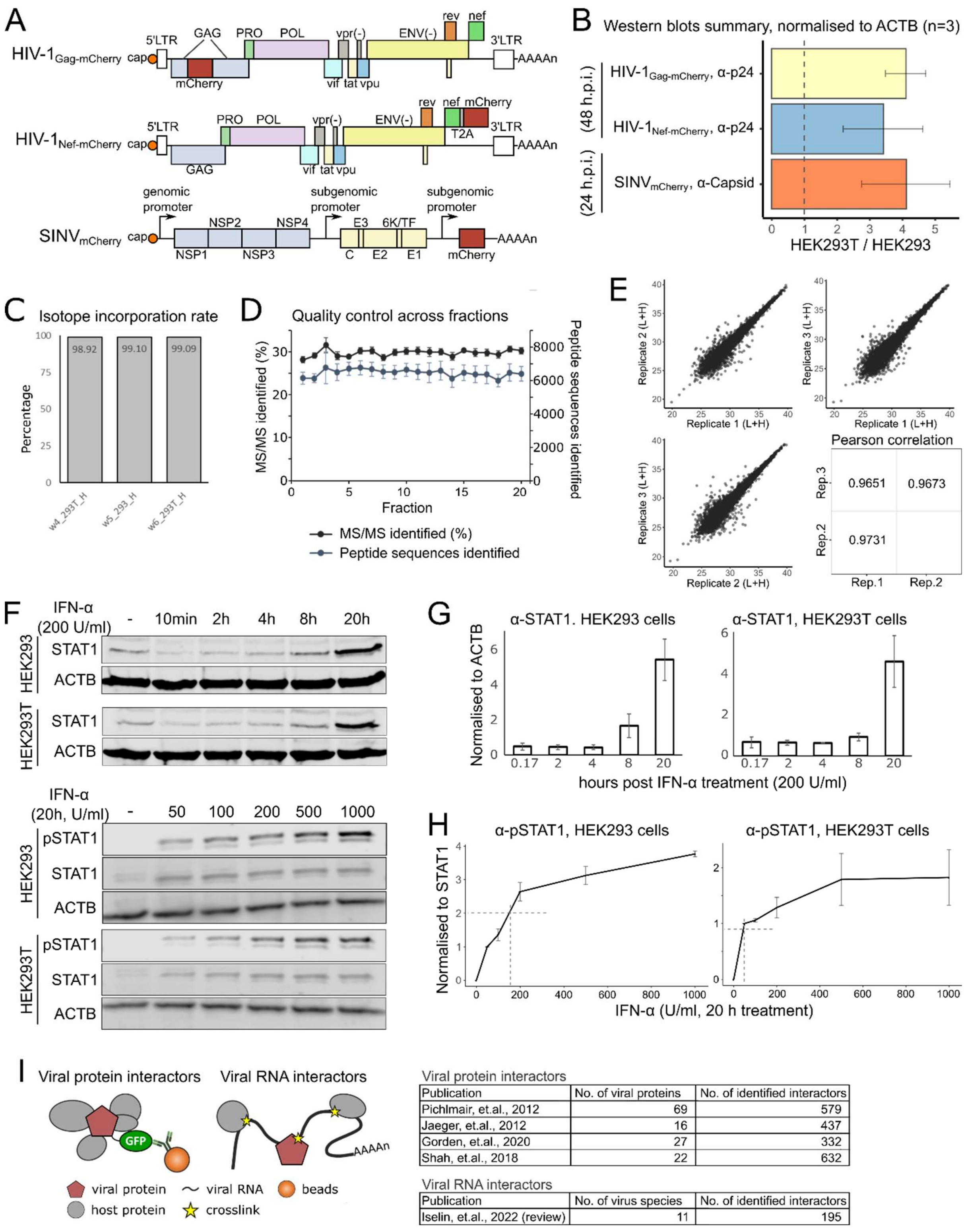


Viral protein interactors

| Publication | No. of viral proteins | No. of identified interactors |
|---|---|---|
| Pichlmair, et.al., 2012 | 69 | 579 |
| Jaeger, et.al., 2012 | 16 | 437 |
| Gorden, et.al., 2020 | 27 | 332 |
| Shah, et.al., 2018 | 22 | 632 |

Viral RNA interactors

| Publication | No. of virus species | No. of identified interactors |
|---|---|---|
| Iselin, et.al., 2022 (review) | 11 | 195 |

**Supplementary figure 1. Deep proteome analysis revealed host processes underlying different permissiveness between HEK293 and HEK293T cells**

A) Diagrams of HIV-1 genome with mCherry insertion in-frame with Gag (HIV-$1_{Gag\text{-}mCherry}$, top) or Nef (HIV-$1_{Nef\text{-}mCherry}$, mid), and SINV genome with mCherry insertion ($SINV_{mCherry}$, bottom).

B) Summary of Western blot experiments comparing viral capsid abundance between HEK293 and HEK293T cells (n=3).

C) Isotope incorporation rates in heavy-isotope labelled SILAC sample, assessed by LC-MSMS.

D) MS/MS identification rates and numbers of peptide sequences identified in each fraction were drawn in line plot for quality control. Error bars represent standard deviation from three replicates.
E) Scatter plots comparing log-transformed intensities between replicates, Pearson correlation coefficients for each contrast pair were shown in corresponding blocks (suppl.tab1).
F) Western blotting with antibodies of ACTB, STAT1, and STAT1-pTyr701 (pSTAT1) from IFN-α treated HEK293 and HEK293T cells. Cells were treated with 200 U/ml of IFN-α for different periods of time (top) or with different dosages of IFN-α for 20 hours (bottom).
G) Barplot summarising ACTB-normalised STAT1 signals from triplicate Western blots as in (F, top).
H) Barplot summarising STAT1-pTyr701 signals normalised to STAT1 from triplicate Western blots as in (F, bottom).
I) Schematic of viral protein/RNA interactors (grey) included in enrichment analysis (left), with tables summarising publications and numbers of interactors determined in each (right).

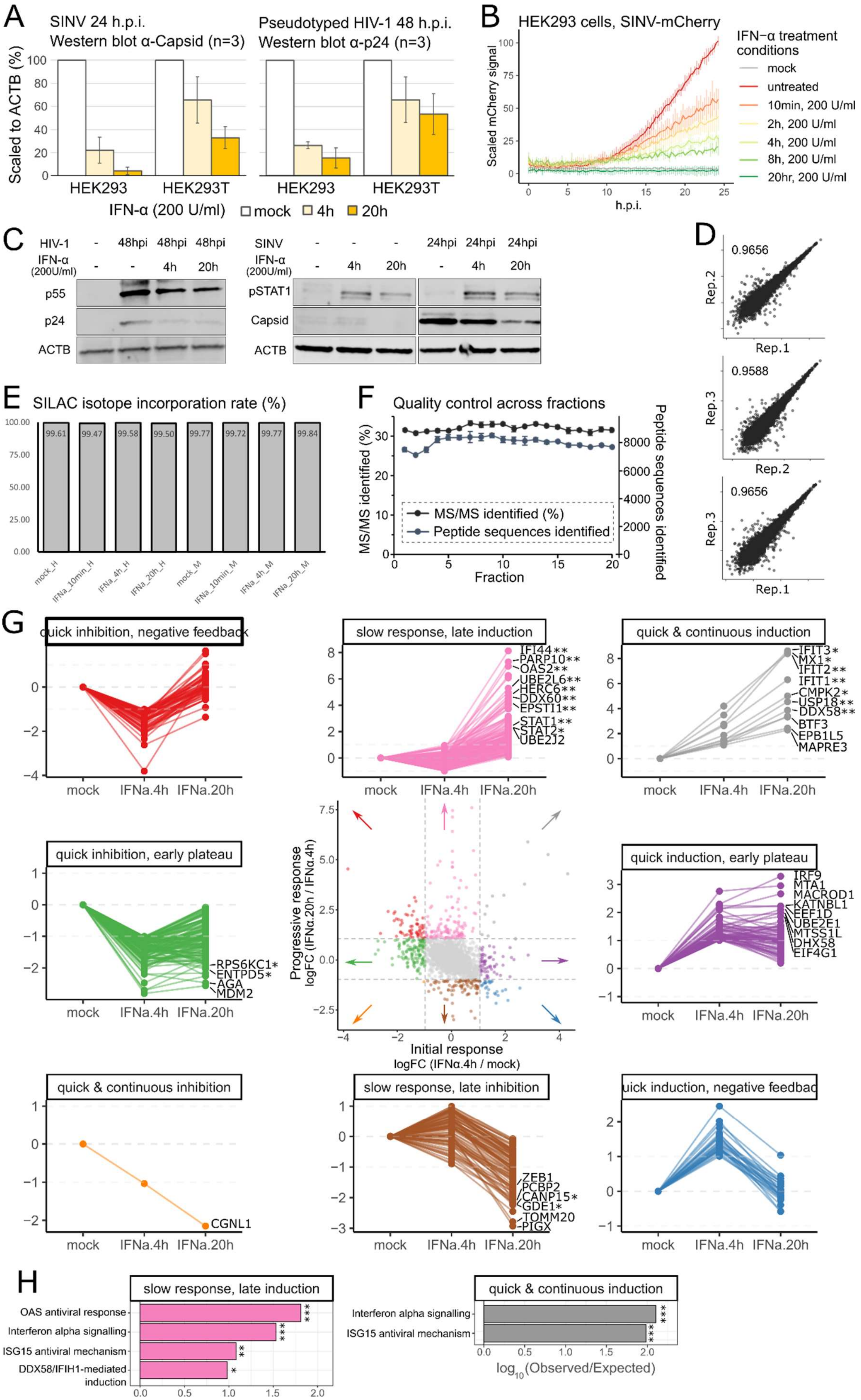

A
SINV 24 h.p.i.
Western blot α-Capsid (n=3)
Pseudotyped HIV-1 48 h.p.i.
Western blot α-p24 (n=3)
Scaled to ACTB (%)
HEK293
HEK293T
IFN-α (200 U/ml)
mock
4h
20h
B
HEK293 cells, SINV-mCherry
Scaled mCherry signal
h.p.i.
IFN−α treatment conditions
mock
untreated
10min, 200 U/ml
2h, 200 U/ml
4h, 200 U/ml
8h, 200 U/ml
20hr, 200 U/ml
C
HIV-1
IFN-α (200U/ml)
48hpi
p55
p24
ACTB
SINV
24hpi
pSTAT1
Capsid
D
0.9656
0.9588
Rep.1
Rep.2
Rep.3
E
SILAC isotope incorporation rate (%)
99.61
99.47
99.58
99.50
99.77
99.72
99.77
99.84
mock_H
IFNa_10min_H
IFNa_4h_H
IFNa_20h_H
mock_M
IFNa_10min_M
IFNa_4h_M
IFNa_20h_M
F
Quality control across fractions
MS/MS identified (%)
Peptide sequences identified
Fraction
G
quick inhibition, negative feedback
slow response, late induction
quick & continuous induction
quick inhibition, early plateau
quick induction, early plateau
quick & continuous inhibition
slow response, late inhibition
uick induction, negative feedbac
mock
IFNa.4h
IFNa.20h
IFI44**
PARP10**
OAS2**
UBE2L6**
HERC6**
DDX60**
EPSTI1**
STAT1**
STAT2*
UBE2J2
IFIT3*
MX1*
IFIT2**
IFIT1**
CMPK2*
USP18**
DDX58**
BTF3
EPB1L5
MAPRE3
RPS6KC1*
ENTPD5*
AGA
MDM2
IRF9
MTA1
MACROD1
KATNBL1
EEF1D
UBE2E1
MTSS1L
DHX58
EIF4G1
Progressive response
logFC (IFNα.20h / IFNα.4h)
Initial response
logFC (IFNα.4h / mock)
CGNL1
ZEB1
PCBP2
CANP15*
GDE1*
TOMM20
PIGX
H
slow response, late induction
OAS antiviral response
Interferon alpha signalling
ISG15 antiviral mechanism
DDX58/IFIH1-mediated induction
quick & continuous induction
log10(Observed/Expected)

**Supplementary figure 2. IFN-α stimulation induce temporal changes in HEK293 cell proteome.**

A) Barplot summarising ACTB-normalised viral capsid protein signals from triplicate Western blots as in (fig.2.F).

B) Infection fitness of $SINV_{mCherry}$ in HEK293 cells treated with 200 U/ml of IFN-α for different periods of time. Fluorescence signals were measured every 15 min in culture condition using a plate reader (top, n=3).

C) Western blottings with antibodies of pSTAT1 and viral capsid proteins. Cells were infected at MOI=1 with $HIV\text{-}1_{Nef\text{-}mCherry}$ or $SINV_{mCherry}$.

D) Isotope incorporation rates in heavy-isotope labelled SILAC sample, assessed by LC-MSMS for IFN-α/mock proteome samples.

E) MS/MS identification rates and numbers of peptide sequences identified in each fraction were drawn in line plot for quality control for IFN-α/mock proteome samples. Error bars represent standard deviation from three replicates.IFN-α/mock proteome samples.

F) Scatter plots comparing log-transformed intensities between replicates for IFN-α/mock proteome samples. Pearson correlation coefficients for each contrast pair were shown in corresponding blocks (suppl.tab2).

G) Scatter plot (centre) comparing log2 fold changes of IFN-α induced initial responses (4 hpt/mock) against progressive responses (20 hpt/4 hpt) for each protein. Temporal changes were divided into 8 groups by cut-off lines at log2 fold change of 1 and visualised in parallel coordinate plots. Top fold changes in each group were text labelled, significant changes were * labelled (*: FDR<0.1, **: FDR<0.01).

H) Enrichment analysis of *Reactome* pathways in groups in (F). P-values of enrichment were labelled in * (*: p<0.05, **: p<0.01, ***: p<0.001).

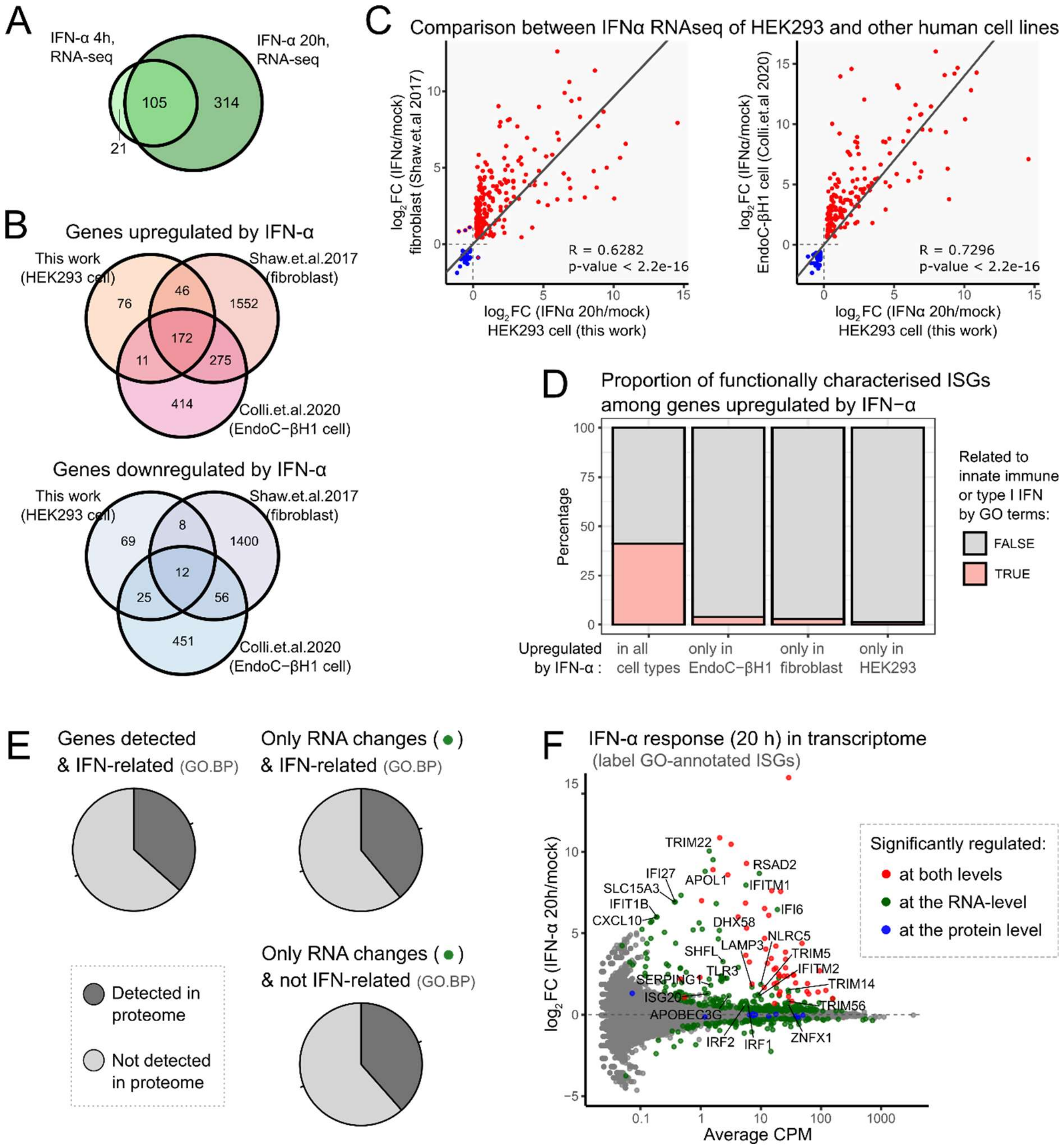


**Supplementary figure 3. IFN-α stimulation induces signature antiviral response in HEK293 cell transcriptome and discordant mRNA-protein changes.**

A) Venn diagrams comparing numbers of IFN-α induced significant changes at 4 hpt and 20 hpt at the RNA level (suppl.tab3).

B) Venn diagrams of RNA-seq results of IFN-α response in different cell lines. Number of genes upregulated (top) or downregulated (bottom) in each cell line and respective publications were compared.

C) Scatter plots comparing log2 fold changes in RNA-seq results between different cell lines. Genes up-/down-regulated in HEK293 cells were filled with red/blue, and genes up-/down-regulated in other cell types were outlined with red/blue.

D) Proportions of GO-BP annotated ISGs among IFN-α upregulated genes identified in different cell lines.

E) Proportion of genes that were detected in proteome among total detected genes, and genes that only showed RNA-level response in RNA-seq.

F) MA plot of IFN-α induced transcriptome change at 20 hpt, with colour labels based on RNA-/protein-levels changes and text labels of GO-BP annotated ISGs.

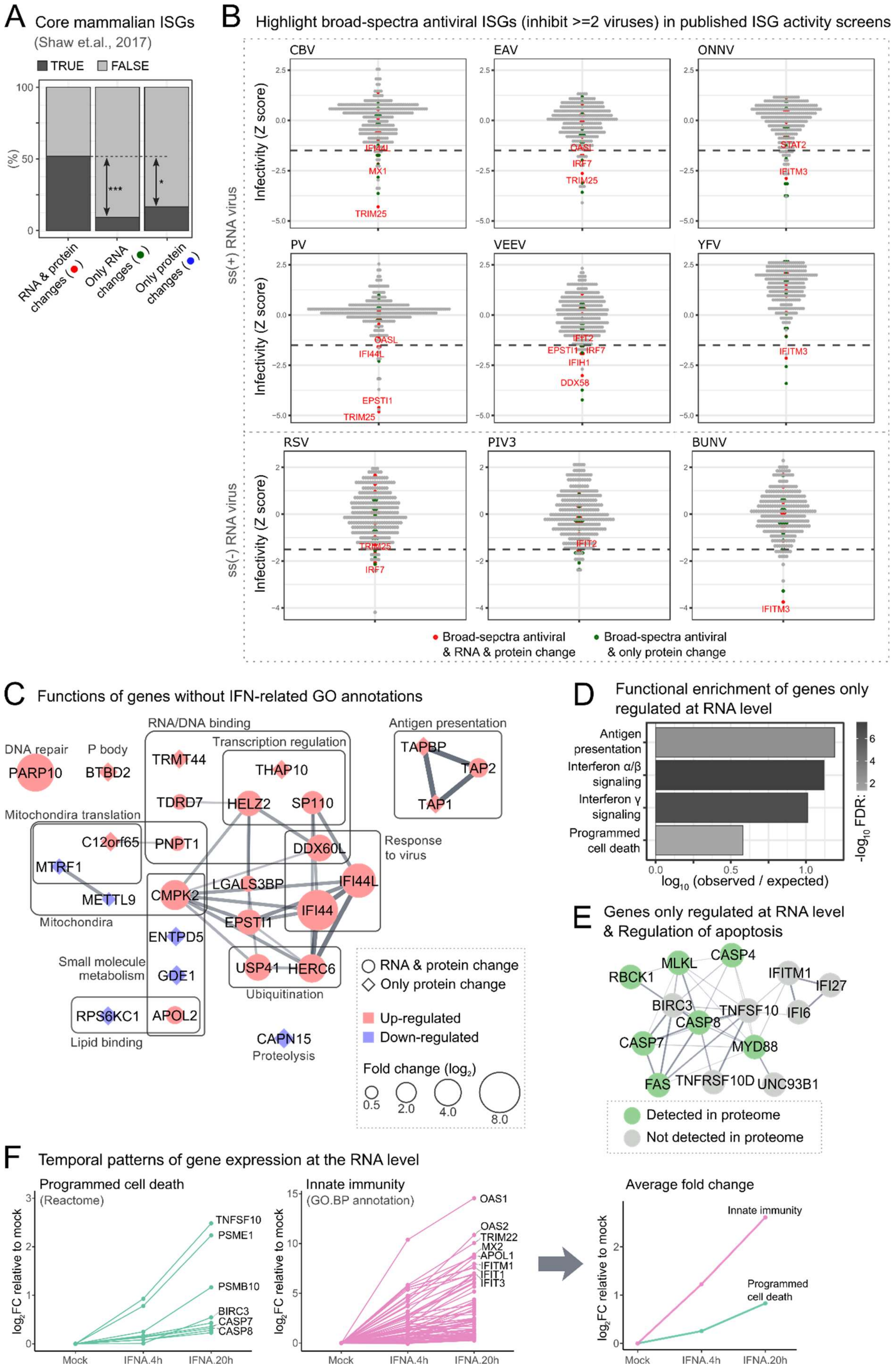

A Core mammalian ISGs
(Shaw et.al., 2017)
TRUE FALSE
(%)
RNA & protein changes
Only RNA changes
Only protein changes
B Highlight broad-spectra antiviral ISGs (inhibit >=2 viruses) in published ISG activity screens
ss(+) RNA virus
ss(-) RNA virus
Infectivity (Z score)
CBV
EAV
ONNV
PV
VEEV
YFV
RSV
PIV3
BUNV
Broad-sepctra antiviral & RNA & protein change
Broad-spectra antiviral & only protein change
C Functions of genes without IFN-related GO annotations
DNA repair
P body
RNA/DNA binding
Transcription regulation
Antigen presentation
Mitochondira translation
Mitochondria
Response to virus
Small molecule metabolism
Lipid binding
Ubiquitination
Proteolysis
RNA & protein change
Only protein change
Up-regulated
Down-regulated
Fold change (log2)
D Functional enrichment of genes only regulated at RNA level
Antigen presentation
Interferon α/β signaling
Interferon γ signaling
Programmed cell death
log10 (observed / expected)
-log10 FDR:
E Genes only regulated at RNA level & Regulation of apoptosis
Detected in proteome
Not detected in proteome
F Temporal patterns of gene expression at the RNA level
Programmed cell death
(Reactome)
Innate immunity
(GO.BP annotation)
Average fold change
log2FC relative to mock
Mock
IFNA.4h
IFNA.20h

**Supplementary figure 4. Antiviral activities and other processes among IFN-α regulated genes in HEK293 cells.**

A) Proportions of core mammalian ISGs (Shaw, Hughes et al. 2017) among genes regulated by IFN-α in HEK293 cells at RNA/protein levels.

B) Dot plots of FACS screening results listed in fig.4B. Genes with broad-spectrum antiviral activities were colour labelled based on IFN-α response at RNA/protein levels in HEK293 cells. ISGs with $Z < -1.5$ for each virus were text labelled.

C) Protein-protein interaction network of genes that have protein-level IFN-α response and have no IFN-related GO-BP annotation.

D) Enrichment analysis of *Reactome* pathways for genes that showed RNA-level, but no protein-level IFN-α response.

E) Protein-protein interaction network of genes involved in regulation of apoptosis, and showed RNA-level, but no protein-level IFN-α response. Genes detected in proteome dataset but determined not significant were green coloured.

F) Parallel coordinate plots showing mRNA expression patterns for genes involved in programmed cell death (left) or innate immunity (mid). Only genes with $FDR < 0.05$ in RNA-seq result were plotted. Overall expression patterns of two processes were compared by respective average fold change (right).

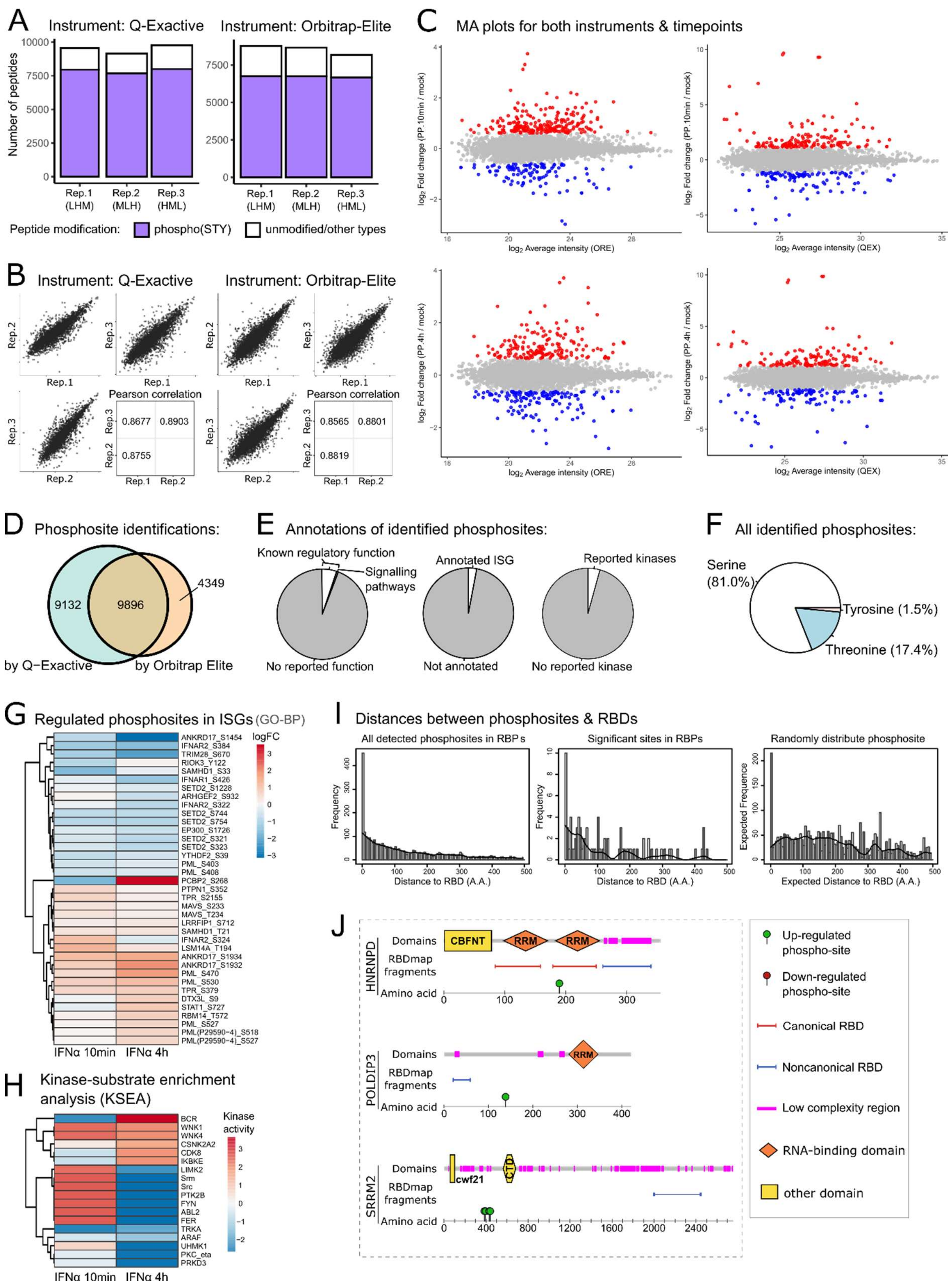


**Supplementary figure 5. Phosphoproteome analysis of HEK293 cells after IFN-α stimulation.**

A) Barplots showing numbers of phosphorylated and non-phosphorylated peptides identified in each SILAC sample from two instrument configurations.

B) Scatter plots comparing log-transformed intensities between replicates, with Pearson correlation coefficients for respective contrast pair. Results from two instrument configurations were displayed separately.

C) MA plots of IFN-α induced phosphoproteomic changes at 10 min and 4 h post treatment obtained with two instrument configurations: Orbitrap-Elite (ORE; left) and Q-Exactive (QEx; right). Phosphosites with FDR < 0.1 in either moderated T statistic or Z statistic were colour labelled and determined as sites of significant changes (suppl.tab4).

D) Venn diagram comparing numbers of phosphosites identified using two instrument configurations.

E) Proportions of functional annotations among all identified phosphosites (left); IFN-related GO-BP annotations among all identified phosphoproteins (mid); kinase annotations among all identified phosphosites (right).

F) Proportions of serine/threonine/tyrosine phosphorylation identified in combined result.

G) Heatmap of IFN-α regulated phosphosites within proteins that have IFN-related GO-BP annotations.

H) Heatmap of kinase-substrate enrichment analysis (KSEA) result (suppl.tab5).

I) Histograms of phosphosites distribution base on their distances (number of amino acids) relative to RBDs.

J) Schematics of RBPs that contains IFN-α regulated phosphosites. “Domains” showed annotations from SMART and Pfam; “RBDmap fragment” showed RNA-interacting peptide fragments identified in proteomics-based interactome analysis (Castello, Fischer et al. 2016).

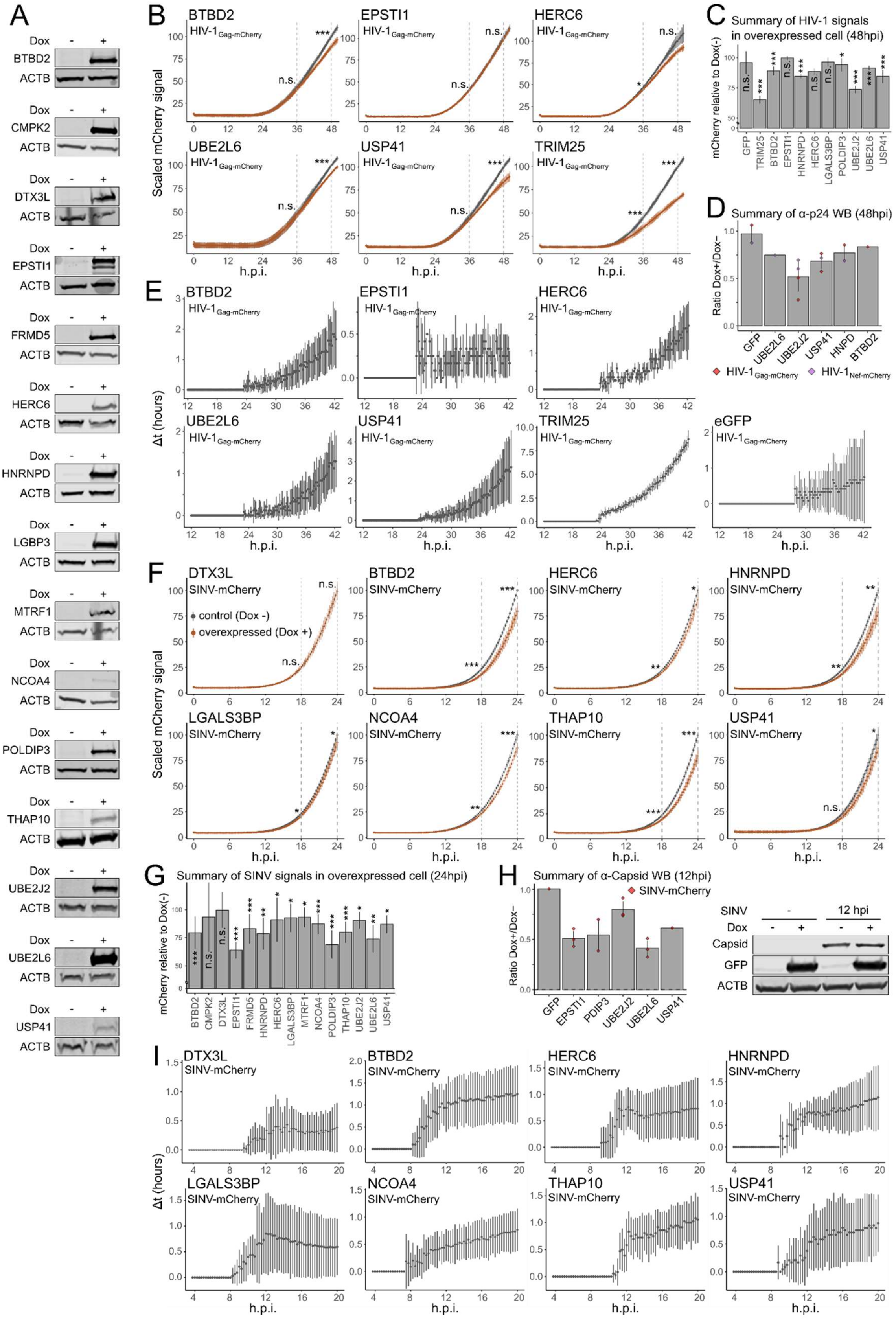


**Supplementary figure 6. Virus-inhibitory effects against SINV and HIV-1 in candidate genes.**

A) Expression candidate genes with eGFP fusion in inducible stable lines verified with Western blottings using antibodies of eGFP and ACTB.

B) Infection fitness of HIV-$1_{Gag\text{-}mCherry}$ in HEK293 inducible stable lines expressing candidate genes with eGFP fusion. Fluorescence signals were measured every 15 min in culture condition using a plate reader (top, n=3).
C) Summary of plate reader results for HIV-$1_{Gag\text{-}mCherry}$ assays as in (B) at 48 hpi.
D) Summary of Western blot experiments that assess virus-inhibitory effects of candidates using HIV-1. Numbers of replicates and type of HIV-1 replicon used were shown in red (HIV-$1_{Gag\text{-}mCherry}$) and purple (HIV-$1_{Nef\text{-}mCherry}$) dots.
E) Delay analysis from 12 to 42 hpi for HIV-1 infected stable lines overexpressing other candidate genes.
F) Infection fitness of $SINV_{mCherry}$ in HEK293 inducible stable lines expressing candidate genes with eGFP fusion. Fluorescence signals were measured every 15 min in culture condition using a plate reader (top, n=3).
G) As in (C) but for assays with $SINV_{mCherry}$ at 24 hpi.
H) As in (D) but for $SINV_{mCherry}$.
I) As in (E) but for SINV infected cells ranging from 4 to 20 hpi.

### References


Castello, A., B. Fischer, C. K. Frese, R. Horos, A. M. Alleaume, S. Foehr, T. Curk, J. Krijgsveld and M. W. Hentze (2016). "Comprehensive Identification of RNA-Binding Domains in Human Cells." Mol Cell **63**(4): 696-710.

Shaw, A. E., J. Hughes, Q. Gu, A. Behdenna, J. B. Singer, T. Dennis, R. J. Orton, M. Varela, R. J. Gifford, S. J. Wilson and M. Palmarini (2017). "Fundamental properties of the mammalian innate immune system revealed by multispecies comparison of type I interferon responses." PLoS Biol **15**(12): e2004086.
